# Pooling size sorted malaise trap fractions to maximise taxon recovery with metabarcoding

**DOI:** 10.1101/2020.06.09.118950

**Authors:** Vasco Elbrecht, Sarah J. Bourlat, Thomas Hörren, Angie Lindner, Adriana Mordente, Niklas W. Noll, Martin Sorg, Vera M.A. Zizka

## Abstract

1. Small and rare specimens can remain undetected when metabarcoding bulk samples with a high size heterogeneity of specimens. This is especially critical for malaise trap samples, where most of the biodiversity is often contributed by small specimens. How to size sort and in which proportions to pool these samples has not been widely explored. We set out to find a size sorting strategy that maximizes taxonomic recovery but remains highly scalable and time efficient.
2. Three 3 malaise trap samples where size sorted into 4 size classes using dry sieving. Each fraction was homogenized and lysed. The corresponding lysates were pooled to simulate samples never sorted, pooled in equal proportions and in 4 different proportions favoring the small size fractions. DNA from the pooled fractions as well as the individual size classes were extracted and metabarcoded using the FwhF2 and Fol-degen-rev primer set. Additionally wet sieving strategies were explored.
3. The small size fractions harbored the highest diversity, and were best represented when pooling in favor of small specimens. Not size sorting a sample leads to a 45-77% decrease in taxon recovery compared to size sorted samples. A size separation into only 2 fractions (below 4 mm and above) can already double taxon recovery compared to not sorting. However, increasing the sequencing depth 3-4 fold can also increase taxon recovery to comparable levels, but remains biased toward biomass rich taxa in the sample.
4. We demonstrate that size fractionizing bulk malaise samples can increase taxon recovery. The most practical approach is wet sieving into two size fractions, and proportional pooling of the lysates in favor of the small size fraction (80-90% volume). However, in large projects with time constraints, increasing sequencing depth can also be an alternative solution.

## Introduction

DNA metabarcoding is a useful approach to characterize arthropod communities. Instead of DNA barcoding individual specimens (Hebert *et al.*, 2003), bulk samples are usually homogenized and DNA extracted. A barcoding marker is then amplified for the whole community and sequenced with high throughput sequencing. This allows for the identification of whole communities (Ji *et al.*, 2013), often with species level resolution given marker choice and reference database completeness (Leray *et al.*, 2019; Weigand *et al.*, 2019). However, DNA metabarcoding is affected by a large number of methodological biases, making it difficult to get accurate specimen counts and often not all taxa in the sample are detected (Elbrecht, Vamos and Meissner, 2017; Piper *et al.*, 2019). While there is a loose correlation between biomass and read abundance (Lamb *et al.*, 2019), metabarcoding results are often skewed by primer bias (Elbrecht and Leese, 2015; Piñol *et al.*, 2015; Krehenwinkel *et al.*, 2017), mitochondrial copy number variation (Braukmann *et al.*, 2019; Deagle *et al.*, 2019), sequencing errors (Singer *et al.*, 2019) and taxa with insufficient biomass may remain undetected (Elbrecht, Peinert and Leese, 2017).

Arthropod bulk samples often show a wide range of specimen sizes (Aylagas *et al.*, 2016). While imagines of the same species often show similar biomass, there can be substantial biomass variation between species and throughout the specimen’s life stages (Stork and Blackburn, 1993; Siemann, Tilman and Haarstad, 1996). Thus, when extracting DNA from bulk samples containing rare or small taxa with little biomass, those might be overshadowed by biomass rich taxa that contribute a large amount of DNA in the extraction. This natural variation in biomass can jeopardize the detection of small and rare specimens (Deagle *et al.*, 2018), especially when taking PCR effects into account (Kelly, Shelton and Gallego, 2019). To maximise taxa detection in bulk samples showing substantial specimen biomass variation (Aylagas *et al.*, 2016), size sorting might be required (Liu *et al.*, 2019). Marine arthropod bulk samples are commonly size fractionated by sieving (Cowart *et al.*, 2015; Leray and Knowlton, 2015; Wangensteen and Turon, 2017; Wangensteen *et al.*, 2018) before metabarcoding. Size sorting was also applied to freshwater macroinvertebrates (Elbrecht, Peinert and Leese, 2017; Carew, Coleman and Hoffmann, 2018; Zizka *et al.*, 2019) and terrestrial arthropods (Ji *et al.*, 2013; Creedy, Ng and Vogler, 2019; Hausmann *et al.*, 2020). While the advantages of size sorting seem logical, the size sorting process can take a substantial amount of time if done manually (Elbrecht, Peinert and Leese, 2017) or even when using semi-automated methods like sieving. Additionally, while size sorting likely increases taxon recovery, it also generates sub samples that can either be pooled or sequenced individually. Regardless of the chosen size sorting and metabarcoding strategy, the amount of laboratory work and costs increases compared to processing a single metabarcoding bulk sample without size sorting. Thus, some authors suggest increasing sequencing depth instead of applying size sorting strategies, especially when size variation is limited (Creedy, Ng and Vogler, 2019) or many of the taxa have differently sized life stages as for freshwater macroinvertebrates (Elbrecht, Peinert and Leese, 2017). In fact, most non-marine studies do metabarcode complete bulk samples without prior size sorting.

In this study, we explore the effects of size sorting malaise trap samples into 4 size fractions, in order to maximise taxon recovery. During DNA extraction, we specifically test the effect of pooling lysates of each size fraction in different proportions. Each size fraction is also extracted and metabarcoded individually, giving us the opportunity to further validate different size sorting scenarios. The goal of this study is to find a standardised and scalable DNA metabarcoding strategy for malaise trap samples, which also captures the diversity of small and rare specimens.

## Material and Methods

### Samples and size sorting

Arthropods were collected on a meadow near Eschweiler, Germany (50.576708N, 6.730254E) using a Townes style malaise trap (Ssymank *et al.*, 2018). The trap was set up in the Natural Park Hohes Venn (calcareous grassland, preserved by grazing animals). The trap was run with 80% denatured ethanol with 1% MEK and samples were collected in 2017 from June 5th to June 26th (sample L1), from July 14th to August 11th (sample L2) and from May 16th to June 5th (sample L3). Samples were dried in an oven (INCU Line ILS 6, VWR, Radnor, PA, USA) at 50°C over 5 days and size sorted with a vibratory sieve shaker (AS 300 control, Retsch, Haan, Germany). Four size fractions were obtained using stacked perforated plate sieves with 8, 4 and 2 mm diameter round holes respectively and a catch basin, each 305mm in diameter (Retsch). Sieving was performed for 5 min with 1.5 mm amplitude. All specimens stuck to the sieves due to electrostatic charging were collected with tweezers. All equipment was thoroughly cleaned with bleach and 5 min UV light exposure between samples.

Size sorting of malaise trap samples with the method outlined above often takes 2-3 hours per sample due to electrostatically charged specimens attaching onto the sieving equipment. Thus, two alternative sieving methods were explored (Figure S1), but not sequenced due to time and financial constraints. For the first method, a “salt shaker” like contraption was built out of two one liter sampling bottles and a 4 mm grid hot glued in a connecting piece, on which both bottles could be attached. Dried specimens could be separated by size by manually shaking the bottle, but were also affected by static charging from the plastic. In a second alternative size sorting approach, a 4 mm grid was placed into a lunch box (with the edges sealed with hot glue). The sample was emptied on top of the grid, and fully immersed in 96% ethanol without prior drying. With a repeated light shaking and pauses of 10 seconds, specimens could be separated into 2 size fractions. The specimens of the small and large size fraction obtained by sieving were dried and all individuals measured in length and weight, to assess the effectiveness of this size sorting approach.

### Tissue homogenisation and lysate pooling

Size fractions were dried overnight and homogenized in 30 ml grinding tubes (Nalgene) with 5 mm diameter steel beads added, using a mixer mill from Retsch (MM400) for 2 min at 30 hz. Each size fraction was transferred to 50 ml falcon tubes or 500 ml VWR bottles and the tissue weight was measured. For tissue lysis, 10 ml ATL lysis buffer (Qiagen, Hilden, Germany) supplemented with 2% Proteinase K (20 mg/ml, VWR) was added per 1 g of tissue. Afterwards, size fractions were shaken overnight at 56 °C using the incubation chamber INCU Line ILS 6 (VWR). After the lysate cooled down, it was centrifuged for 5 min at 4149 g in a VWR centrifuge (Mega Star 1.6). The lysates of the 4 size fractions were pooled in different proportions (Figure 1 & Table S1). Since the amount of extraction buffer added is proportional to the amount of tissue in the respective size fraction, we assumed the DNA concentration in each lysate to be the same. Thus, pooling lysate volumes of the 4 size fractions proportionally to the tissue weight, simulates a full malaise sample as if it was not size sorted. The other proportions used for pooling the size fractions test if equal pooling or pooling in favor of the small fractions increases taxon recovery compared to the unsorted malaise trap sample. DNA was extracted from the lysate using a Qiagen DNeasy 96 well Blood & Tissue Kit (Qiagen, Hilden, Germany). Finally, DNA quality was examined via gel electrophoresis on a 0.5% agarose gel (VWR) with 0.1% GelRed (10000x, Biotium) and 1x TBE (VWR). 1µl of GeneRuler 100bp plus DNA ladder (Thermo Scientific) and 3µl of PCR product were loaded into the gel. The plate map with sample locations is available as supporting information (Figure S2), with 27 wells being used for primer tests as part of a separate project and 9 negative controls in which only lysis buffer and proteinase K without tissue were applied for extraction with the DNeasy 96 well Blood & Tissue Kit. The eluates obtained from these negative controls were also used for the amplicon and indexing PCR.

**Figure 1:**
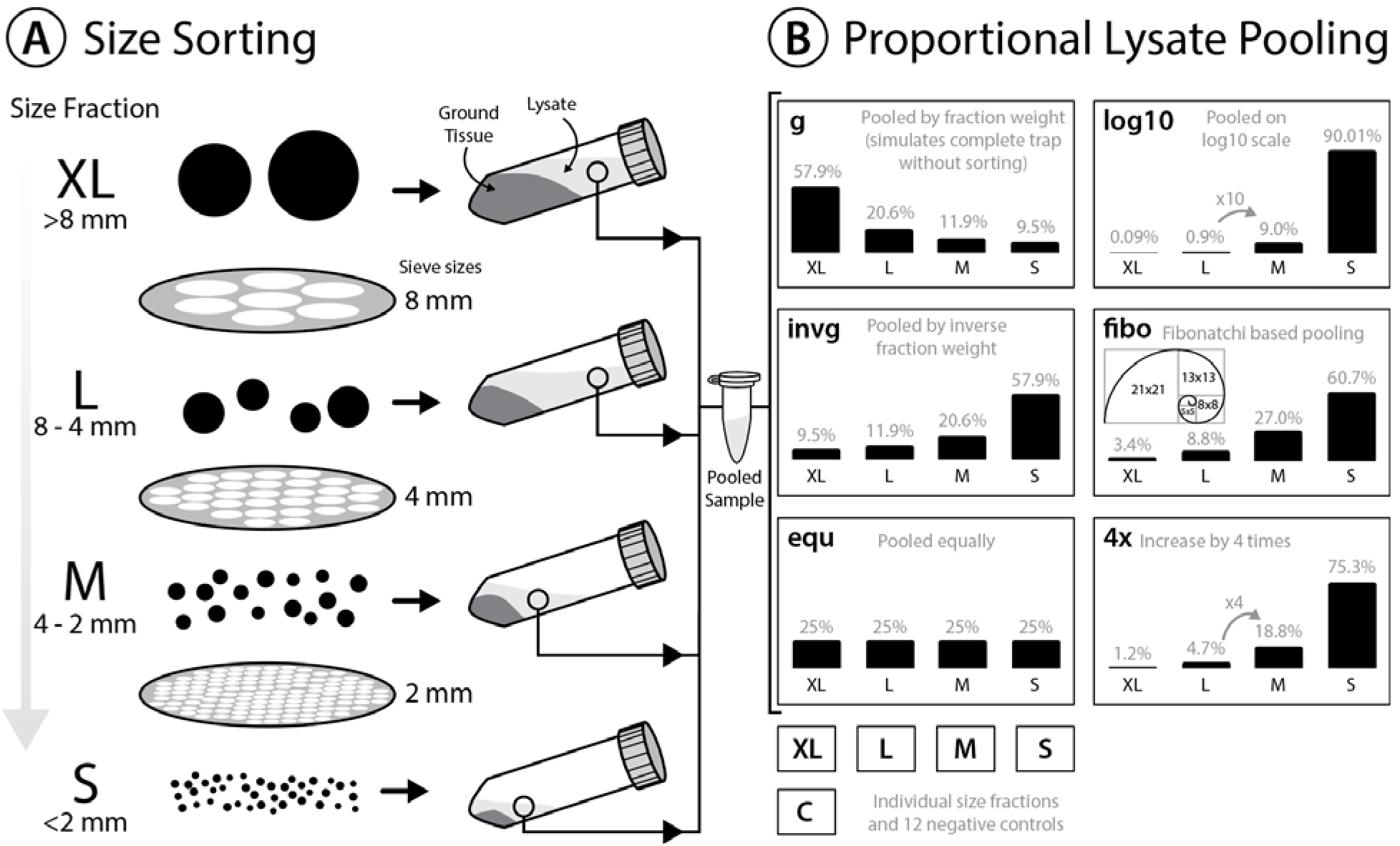
Experimental design to test the effectiveness of size sorting malaise trap samples and pooling the fractions in different proportions. **A:** Dried specimens are sieved into 4 different size classes, homogenized and lysed with a buffer amount corresponding to specimen weight in the respective fraction. **B:** Lysate is incubated, and pooled in different amounts. Gray numbers above the bars indicate the relative proportions (%). All 6 lysate pools were generated in duplicate, and each individual size fraction (S, L, M, XL) was also extracted in duplicate, as well as 9 negative controls included.

### DNA metabarcoding

Sample metabarcoding was carried out using a 2 step approach, with standard illumina Nextera primers used for dual tagging in PCR 2 (index PCR). PCR 1 (amplicon PCR) was carried out in a 96 well plate using reactions of 25 µL with 1 µL DNA (stock undiluted), 0.2 µM of each fwhF2 forward primer (Vamos, Elbrecht and Leese, 2017) and Fol-degen-rev reverse primer (Yu *et al.*, 2012), 12.5 µL PCR Multiplex Plus buffer (Qiagen, Hilden, Germany) and 10.5µl ddH_2_O (Sigma-Aldrich). Primers included a universal Nextera tail to facilitate sample tagging in PCR 2. The thermal cycler 2720 from AB was used and set with the initial denaturation at 95 °C for 5 min; 30 cycles of: 30 sec at 95 °C, 30 sec at 50 °C and 50 sec at 72 °C; and a final extension of 5 min at 72 °C. 1 µL of PCR product was used, without cleanup (Elbrecht and Steinke, 2019), as template for the second PCR. In the second PCR, 12 times 8 Nextera (Illumina) based primer combinations were used, to attach 8 bp dual indexes to individual samples as well as the tails needed for Illumina sequencing. The following Nextera XT Index Kit v2 adapters i5 (Index 2, forward) and i7 (Index 1, reverse) were used: S503, S505, S506, S507, S508, S510, S511, S517 and iN702, iN703, iN704, iN705, iN711, iN712, iN714, iN715, iN718, iN723, iN724, iN727. The 25µl reactions in the second PCR were composed of 1 µL PCR1 (undiluted, no prior cleanup), 0.4 µM of each primer (Nextera, Illumina, San Diego, CA, USA), 12.5 µL PCR Multiplex Plus buffer (Qiagen, Hilden, Germany) and 10.5µl ddH_2_O. The same PCR cycling conditions as in the first PCR step were used. PCR success was checked on a 0.7% and 1% agarose gel for the first and second PCR, respectively. Gels were composed as described above for the DNA quality check. PCR products were normalized using SequalPrep Normalization Plates (Thermo Fisher Scientific, MA, USA; (Harris *et al.*, 2010) to obtain 25ng product per reaction according to manufacturer protocols. Normalization success was verified optically on a 1% agarose gel also as described above. 10 µL of each normalized sample library was pooled, and the final library was cleaned up twice using left sided size selection with 0.76x and 0.65x SPRIselect (Beckman Coulter, CA, USA), to remove primer dimers from the negative controls. The library concentration (2.01ng/µl) was measured using the QuantiFluor dsDNA system (E2670, Promega) and a Quantus fluorometer (Promega, Madison, WI, USA). The cleanup success was verified by running samples on a Fragment Analyzer (Agilent Technologies, Santa Clara, CA, USA) using the HS NGS Fragment Kit. The Fragment Analyzer data was evaluated using the software PROSize 2.0. Sequencing was carried out by Starseq (Mainz, Germany) using a 600 cycle Illumina MiSeq Reagent Kit v3 and 5% PhiX spike in.

### Bioinformatic processing

Raw sequence data was delivered in blc format, and converted to fastq files using bcl2fastq v2.20.0.422 using the option `--create-fastq-for-index-reads` and sequence quality verified using FastQC v0.11.9. Since there were No call issues with the Sequencing run, the base calls for each flow cell tile in each cycle were plotted and evaluated for problems using the `PlotTiles` function in JAMP v0.XXX (github.com/VascoElbrecht/JAMP). No call events were limited to the primer binding regions. Sequences where demultiplexed using deML using exact matching (github commit dd87669, (Renaud *et al.*, 2015)) and the demultiplexing table provided in Scripts S1. Sequence data was processed using the JAMP pipeline that mostly relies on Vsearch v2.14.2 (Rognes *et al.*, 2016) and Cutadapt v2.8 with allowing up to 30% mismatches in the primer binding region (Martin, 2011). The used scripts are available in Scripts S1. Paired end merging of read 1 and read 2 was done using vsearch with `fastq_maxdiffpct = 25` to maximise the amount of reads merged. Primer sequences were removed from each sample using Cutadapt, allowing for 30% mismatches in the binding region, while retaining only sequences where both primers were successfully trimmed. Cutadapt was also used to remove sequences below 303 bp and above 323 bp length. Poor quality sequences with an expected error score (Edgar and Flyvbjerg, 2015) above 1 or containing N base calls where removed using vsearch. All samples were pooled and dereplicated (`minsize=2`) and clustered into OTUs using vsearch with `cluster_smallmem` and 3% similarity, followed by denovo chimera removal. Individual samples were dereplicated (retaining singletons) and mapped against the OTUs to generate an OTU table, using `usearch_global` with `maxrejects=256` and `maxaccepts=16`. Potential false OTUs were detected using lulu version from August 2018 (Frøslev *et al.*, 2017). Before collapsing flagged OTUs, taxonomy was assigned using https://www.gbif.org/tools/sequence-id and only flagged OTUs that share the same taxonomy merged. Data was further cleaned up by multiplying the maximum read number of each OTU in the negative controls, and subtracting it from all other samples (Elbrecht and Steinke, 2019). Additionally, all read counts below 0.01% abundance in at least one sample and both replicates were discarded. Filtering all samples at 0.01% relative abundance ensures comparisons are done at the same sequencing depth, while reducing stochastic effects introduced by repeated rarefaction of the data. Filtering at 0.01% abundance is similar to a sequencing depth of 10.000 sequences per sample. Additionally, since all reads which are not present in both replicates with 0.01% abundance were discarded, the 2 replicates were combined for statistical analysis. Since the gbif database is clustered at 99% similarity, BOLDigger v1.1.3 (https://github.com/DominikBuchner/BOLDigger) was used to assign taxonomy to the OTU subset to improve taxonomic matching (setting ‘JAMP pipeline’).

## Results

Sieving dried specimens into 4 size fractions using the “sieving tower” (Figure S1 A) did sort specimens by size, but took 2-3 hours per sample as statically charged specimens and broken off tissue pieces had to be collected from the sieve surfaces. Thus an alternative “shaker” design was tested, but was affected by the same static charging issue. The wet sieving strategy however did not show this issue and allowed size separation in under 10 minutes, by alternating between gently shaking and letting the specimens sink down, with small specimens falling through the 4 mm mesh. The wet sieving did lead to a clear size fractionation, with one order of magnitude difference in average specimen weight between the small and large size fraction (Figure S1 B). However, for sequencing, only the 3 dried samples were used, as the alternative size sorting strategies were explored afterwards.

The MiSeq run generated 21.911.822 PE reads which are available as raw data as well as demultiplexed samples under the NCBI SRA accession PRJNA625408. No call events were observed on several tiles in read 2, leading to N being inserted at positions 16, 17 and 25. A total of 20 out of 38 tiles were affected by one or more No Call events, but as affected areas were mostly removed by the primer trimming, this had only a minor influence on the analysis. Average sequencing depth per sample was 249.423 reads (SD = 92.075) with the lowest sample containing 146.344 reads. In bioinformatics processing an average 23.40% (SD = 4.16%) of reads were discarded. Additional run statistics as well as raw and filtered OTU tables are available as supporting information in Scripts S1 and Table S2. After all filtering steps were applied, 99% of the 2197 OTUs were assigned to the phylum Arthropoda.

To assign a specimen size to each OTU, relative read abundance of all 3 samples was combined for each size class, and the size class with the highest abundance selected as a specimen size estimate. Three quarters of OTUs were present in only one size fraction (Figure S3). Thus most OTUs could be clearly assigned to a size class using this strategy, with a few exceptions in the large and medium size category (e.g. OTU1 *Myrmica ruginodis* and OTU11 *Sphaerophoria*). Nevertheless, the most abundant size category was assigned, even when an OTU was found in high abundance in both samples. For some low abundant OTUs, no reads were found in any of the size classes, but in the pooled samples (this is the case for 9.38% of OTUs). Thus, the respective OTUs had no size class assigned. Most OTUs were assigned to smaller size classes, with 41.51% in S, 24.53% in M, 19.21% in L and 5.37% in XL (Table S1). The individual size fractions (S, M, L and XL) recovered mostly reads from their respective size class (Figure 2), with only a few exceptions like OTU1 (*Myrmica ruginodis*). Here both the large and medium size fraction a large amount of reads (43.2% and 55.7%). Thus, for OTU1 the size class medium is assigned, despite the large size fraction also containing a substantial amount of reads. The total number of taxa recovered was the highest in the smaller size fractions, and only below 100 OTUs recovered in the extra large size fractions (Figure 2). When pooling the lysates based on dry size fraction weight (g) to simulate a completely unsorted malaise sample, the samples are dominated by large specimens. Compared to the other samples pooled in various proportions, the unsorted malaise traps showed the weakest OTU recovery, followed by pooling all 4 lysates in equal proportions (equ). The other four methods of pooling (invg, fibo, 4x and log) did all recover more OTUs, with equally high numbers detected across all 3 samples. The logarithmic pooling did recover the highest proportion of small specimens, while showing reduced recovery of large specimens compared to the other methods.

**Figure 2:**
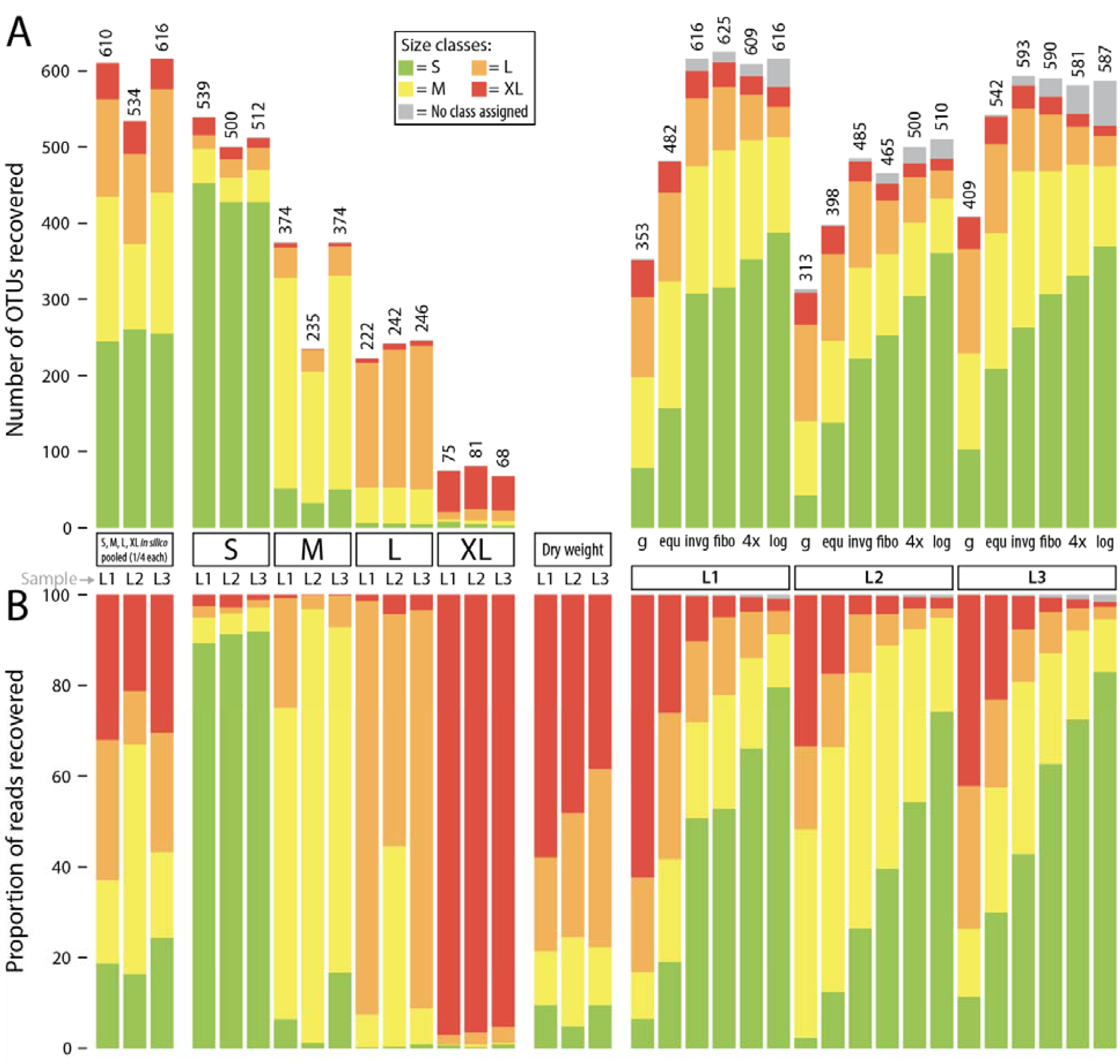
Detected OTUs across different samples (L1,L2 and L3). OTUs are assigned to size classes using the individually sequenced size fractions (S, M, L, XL). Some low abundant OTUs were not recovered in individual size fractions, but in pooled samples. Thus a size class was not assigned (gray bars). The 4 individual fractions were also pooled at 1/4 sequencing depth (each filtered with 0.04% threshold) to simulate equally pooling the samples *in silico*. **A:** Barplots showing the number of OTUs recovered from individual size fractions as well as pooled samples. **B**: Barplot showing relative read abundance across samples (L1,L2 and L3). The “Dry weight” bars are showing the expected distribution of size classes based on relative abundance of dry weight of each size fraction, allowing a comparison to the proportionally pooled samples (g) that simulate a complete unsorted malaise trap sample.

When pooling the 4 individually sequenced size fractions at 1/4 sequencing depth of the lysates pooled in equal proportions (equ), the individually sequenced size fractions did recover an average of 26.56% more OTUs than the equally pooled lysates (Figure 2). However, the numbers of OTUs recovered from the *in silico* pool of individually sequenced size fractions were comparable to OTU counts recovered by lysates pooled in favour of small size fractions. The read proportions of *in silico* equally pooled samples were similar to the proportions of the pooled lysates (equ).

In the rarefaction analysis, the samples pooled in favor of small size fractions (invg, fibo,4x, log) did recover a similar amount of OTUs at around 15.000 reads sequencing depth, as the unsorted sample (g) at 50.000 reads (Figure 3). This approximately represents a factor of 3-4x reduction of sequencing depth, compared to unsorted samples. While the number of OTUs might be similar in this comparison, unsorted samples (g) are more biased towards large specimens, while the samples pooled in favor of small specimens recover less large specimens (Figure 2). Thus, while the number of OTUs recovered from the unsorted samples at 3-4x sequencing depth might be similar to these size sorted samples, there are still differences in OTU composition between differently pooled samples.

**Figure 3:**
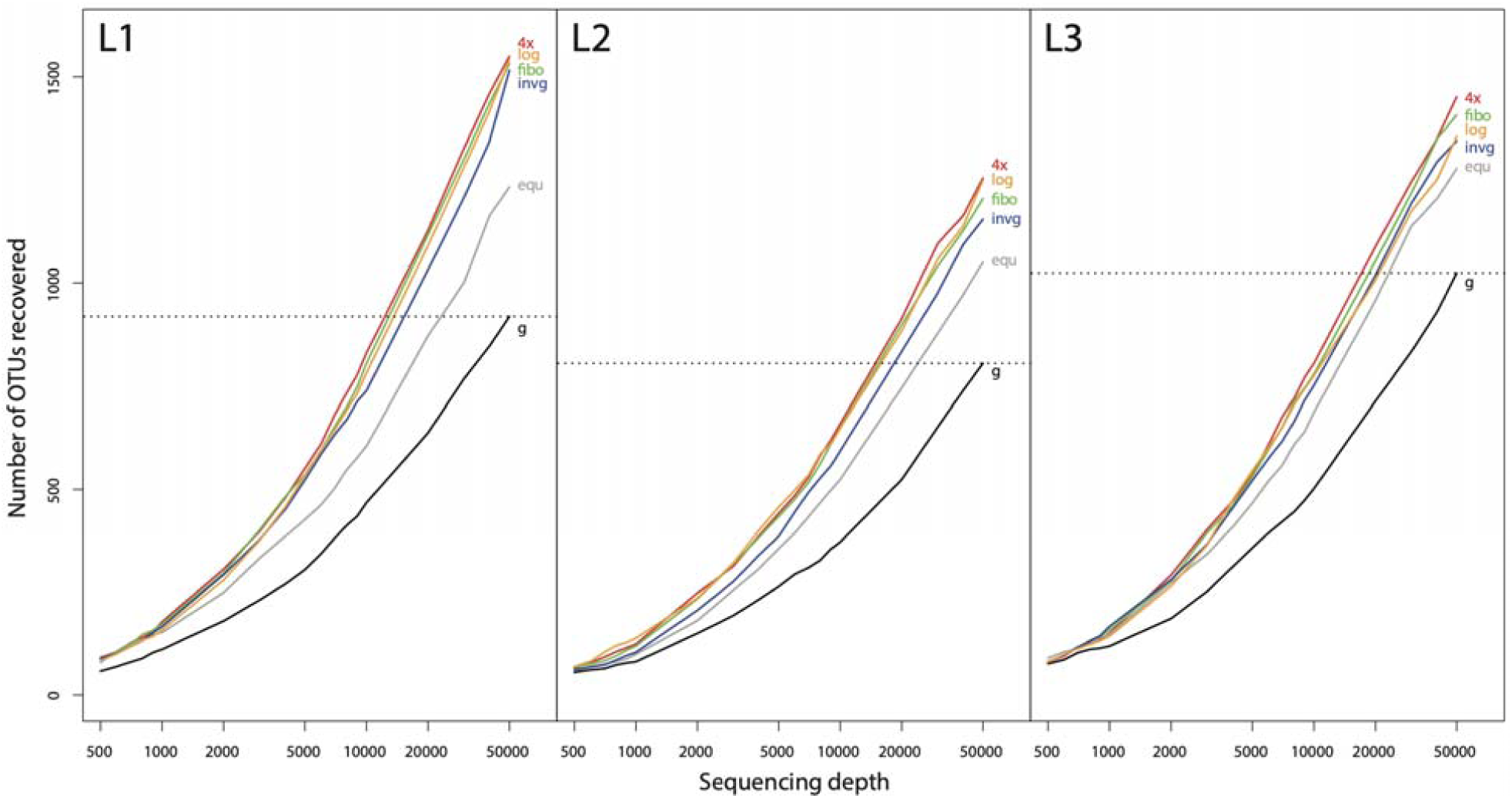
Rarefaction curves at different sequencing depths for all 3 samples (L1,L2 and L3) and lysates pooled in different proportions. The taxon recovery of the unsorted malaise sample (g) at 50.000 reads sequencing depth is used to estimate the increase in sequencing depth needed to recover a similar amount of OTUs than in samples that are size sorted and pooled in favor of smaller size fractions. The OTU counts in this rarefaction analysis are slightly higher than in our other analyses, since replicates were pooled directly, without removing reads that are not in both replicates and without discarding OTUs or reads below 0.01% abundance.

To test if separation of a sample in only 2 size fractions would be sufficient to obtain optimal taxon recovery, individual size fractions S+M and L+XL were combined and pooled *in silico* in different ratios (Figure S4). The recovery of OTUs was highest when adding 10-20% of the large fraction to the small fraction. The overall OTU recovery in the *in silico* pooled samples did in fact exceed the number of OTUs recovered by the metabarcoded lysate pools (invg, fibo, 4x, log) and sample L1 and L3 by up to 10-20%.

## Discussion

Previous studies have demonstrated that size sorting before metabarcoding can increase taxon recovery (Elbrecht, Peinert and Leese, 2017; Creedy, Ng and Vogler, 2019), especially when dealing with bulk samples containing specimens with a wide biomass variability. We were able to confirm these trends for malaise trap samples. The small and medium size fractions contained the majority of the diversity in each of the three samples, consistent with studies from marine benthos (Wangensteen *et al.*, 2018). When pooling four size fractions in different proportions, the fractions simulating a complete malaise trap without size sorting was dominated by large specimens, showing consistently the lowest recovery of OTUs. Interestingly, all four ways of pooling in favour of the smaller fractions (inverse weight, logarithmically, fibonacci based and four fold increase) all recovered a similar number of OTUs. This result was somewhat unexpected, as for example logarithmic pooling does strongly favour small fractions which would make large size fractions severely underrepresented. However, this effect was not as pronounced, since there is some OTU sharing across the individually sequenced size classes. This might be caused by small specimens being attached to larger specimens or contained as prey in their gut. Additionally, legs and other tissue pieces can easily break off when rigorously sieving the dry specimens, and thus tissue from large specimens can end up in small size fractions. This might be advantageous, enabling detection of larger specimens when sequencing smaller size fractions. But small pieces of tissue might not break off from all species with the same chance, thus the large taxa found in the small fractions might not be representative of the real diversity in the samples. Of course, different size fractions should be pooled in the same proportions to ensure comparability across the samples. The amount of small fraction used in the DNA extraction will depend on the goals of the study, and what size fractions are mostly of interest, given that the number of reads and OTUs of differently sized specimens is dependent on the pooling process.

Now as we have established that size sorting can increase taxonomic recovery in malaise trap samples, the question is how to size sort and process arthropod bulk samples efficiently. Only if the size sorting process is efficient, and the amount of laboratory work does not increase substantially, size sorting can be used in larger scale metabarcoding projects. When the amount of samples is limited and the variation in specimen biomass not substantial, biomass introduced biases can be limited (Creedy, Ng and Vogler, 2019) or can be minimized by only including a small amount of biomass (e.g. one leg) from the large specimens (Ji *et al.*, 2013; Carew, Coleman and Hoffmann, 2018). While manual sorting and processing of specimens can be a good strategy to reduce biomass bias, it is quite work intensive and difficult to scale or standardise. Here a more automated approach would be desirable. However automated sieving approaches also come with their own challenges. For example, with the dry sieving approach taken in this study, we ran into major methodological limitations, as it took 2-3 hours per sample picking individual electrostatically charged specimens from the sieving equipment. Here we pioneered an alternative using wet sieving, completely immersing the specimens and sieve in ethanol, contained in a lunch box. This approach can likely be automated using incubation shakers, reducing the hands on time and increasing scalability of this size sorting approach by an order of magnitude. Additionally, since only gentle shaking is used with wet sieving, specimens are less damaged and might be used for morphological identification if non destructive metabarcoding is applied (using ethanol (Hajibabaei *et al.*, 2012) or lysis buffer protocols (Nielsen *et al.*, 2019)). However, non destructive methods can introduce additional biases to the metabarcoding process, in some cases detecting substantially less taxa than with morphology or homogenizing bulk samples (Erdozain *et al.*, 2019; Marquina *et al.*, 2019). To generate multiple size fractions, several sieves can be placed over each other. However our *in silico* evaluations show that size sorting with one single 4 mm sieve into a small and large size fraction represents a good trade off between taxon recovery and laboratory workload. Pooling the large and small size fraction in the proportion 1:10 to 2:10 did show the highest taxon recovery in our in silico test, but this should be confirmed experimentally, since *in silico* comparisons are limited in reliability. For example, when *in silico* pooling 1/4 of each of the individually sequenced size fractions, more taxa were recovered compared to actually pooling the 4 size fractions equally and then sequencing them. Thus, the taxon recovery for the *in silico* estimates are likely to be slightly overrepresented. This can likely be explained by primer bias effects (Elbrecht and Leese, 2015; Piñol *et al.*, 2015) that can become quite pronounced when pooling size fractions together. For example, if in the large size fraction a specific taxon is very well amplified, it will make up a large amount of the reads in the pooled sample, decreasing the sequencing depth of all other specimens. When however individual size fractions are sequenced, these negative primer effects are limited to each individual fraction, increasing overall taxon recovery. For the same reasons, some authors recommend sorting metabarcoding bulk samples taxonomically and sequencing each individual taxonomic group (Morinière *et al.*, 2016; Beentjes *et al.*, 2019). A taxonomic sorting step is however very impractical for metabarcoding, increasing processing time per sample substantially, while not substantially increasing taxon recovery given that most arthropod primer binding sites show the same pattern of variability (Elbrecht and Leese, 2017; Elbrecht *et al.*, 2019).

Some studies did sequence each individual size fraction (Cowart *et al.*, 2015; Wangensteen *et al.*, 2018). This can be advantageous, as it provides information about the community composition for each size class, which could be useful depending on the study context. This can be the case particularly when larval stages are collected, which can show a range of specimen sizes that will also be reflected in individually sequenced size fractions (Elbrecht, Peinert and Leese, 2017). Arthropods collected in a malaise trap are however often adult insects, that are less variable in specimen size than larval stages throughout their life history. Thus, in most cases, the sequencing of individual size fractions will not be needed when lysates or extracted DNA is pooled proportionally to reflect the small size fractions sufficiently. Size sorting and then pooling lysates of each fraction before the DNA extraction can be especially advantageous, as the metabarcoding workflow can stay in a 96 well plate format (Elbrecht and Steinke, 2019). Additionally, sample processing costs and time can be substantially reduced by processing pooled lysates, compared to sequencing each size fraction individually.

While we confirm that size sorting can increase taxon recovery when metabarcoding malaise traps samples, the implementation (e.g. number of size fractions, proportions of pooling) will depend on the aim of the study. In some cases, size sorting might be skipped all together, as a sufficient signal can be achieved with sequencing depth, especially when specimen size variability is limited (Creedy, Ng and Vogler, 2019). While Creedy et al. 2019 do not give exact numbers on sequencing depth effects, we estimate that with a sequencing depth increase of 3-4 fold, similar numbers of OTUs will be recovered from an unsorted sample as with size sorting in 4 fractions and pooling in favor of small specimens. However, by increasing the sequencing depth, the metabarcoding results will still be dominated by biomass rich taxa, with many less biomass rich taxa remaining undetected as a consequence. Thus, in cases where low abundant and small taxa are relevant, sequencing depth has to be either increased further or the sample has to be size sorted to increase representation of small taxa. Given that size sorting can be done in around 15 minutes using the wet sieving method demonstrated here, size sorting seems like a worthwhile step for malaise trap samples. Further malaise trap sampling and additional size sorting experiments could be useful to further quantify the effects of large biomass rich specimens on the detection of small or rare specimens, helping to develop standardised protocols for biomonitoring of flying arthropods.

## Conclusions

We demonstrate that size sorting of malaise trap samples can increase recovery of small specimens. Pooling lysates in the extraction process can be an efficient way of improving representation of small taxa, while not increasing laboratory work and costs substantially, as would be the case with sequencing each fraction individually. *In silico* tests show that only two size fractions (4mm sieve) pooled 1:10 to 2:10 can already increase taxon recovery substantially. Increasing sequencing depth 3-4x can also be an alternative to size sorting, but might still under represent small and rare taxa. Given that wet sieving of specimens in ethanol is efficient and quick, size sorting can be considered an effective approach to increase taxonomic recovery of malaise trap samples with DNA metabarcoding. If size sorting is needed depends however, on the specimen size variation in the samples as well as the research goals and taxonomic recovery required.

## Supporting information

Scripts S1

## Supporting information

Fig S1: Comparison of different size sorting strategies.

Fig S2: Plate map showing position of samples as well as illumina indexing for the different samples.

Fig S3: Specimen size overlap in individually sequenced size fractions.

Fig S4: Heatmap showing OTUs recovered for each sample, and size class assigned to each OTU.

Fig S5: *In silico* pooling of individually sequenced size fractions S & M with L & XL in different proportions.

Scripts S1: JAMP scripts used for data processing and R scripts to generate figures.

Tab S1: Table showing dry weights of each size fraction and amount of lysate pooled for each sample.

Tab S2: Raw OTU table and OTU table used for analysis, with pooled replicates, low abundant OTUs removed and similar OTUs collapsed with LULU.

## Acknowledgements

We would like to thank the entomological society Krefeld for providing the malaise trap samples. Claudia Etzbauer for laboratory infrastructure and supplying us with additional ATL buffer. Thomas Creedy, for helpful discussions and/or feedback to the manuscript. Sandra Kukowka for running the library on the fragment analyzer.

## Author contributions

T.H. and M.S. did provide the malaise trap samples. V.E. and A.L. developed the concept and carried laboratory work. A.M. did measure and weight size sorted specimens. V.E. analysed the data with help of N.W.N., V.Z. and A.M.. V.E. did write the manuscript with help of all authors.

## Data Accessibility statement

Raw sequence used in this study data is available in the NCBI short read archive under the accession number PRJNA625408.

