## Supplementary material for "Pooling size sorted malaise trap fractions to maximise taxon recovery with metabarcoding": Figure S1

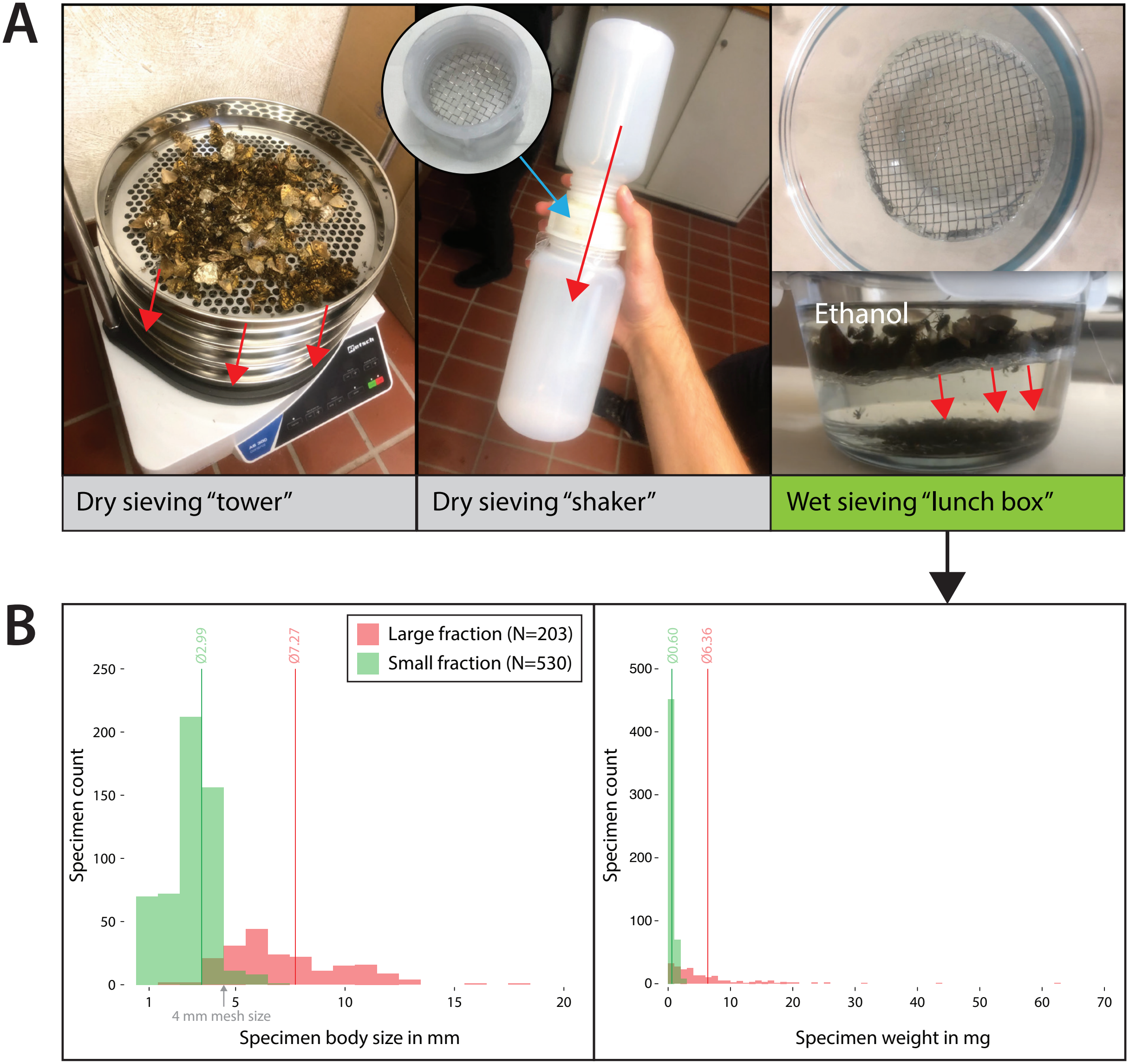


**Fig. S1:** Overview of tested size sorting strategies. **A:** Different dry sieving strategies ("tower" and "shaker", time consuming due to electrostatic charging of specimens) and wet sieving strategy in ethanol using a lunch box and a 4 mm metal mesh. **B:** Histogram showing the size and weight differences of one additional malaise sample from Vlotho (52.122705N, 8.785186E) wet sieved into two size fractions using the "lunch box" design. Specimen body size measured without antennae or legs. Sample was collected in 2017 from September 19th to October 3rd.
