## Supplementary material for "Pooling size sorted malaise trap fractions to maximise taxon recovery with metabarcoding": Figure S2

| index 1: \ index 2: | S517<br>H | S511<br>G | S510<br>F | S508<br>E | S507<br>D | S506<br>C | S505<br>B | S503<br>A |
|---|---|---|---|---|---|---|---|---|
| N702 1 | D<br>C01 | D<br>L1_MA | D<br>L1_XLB | D<br>L2_fiboA | D<br>L2_equB | D<br>L2_SB | D<br>L3_XLA | D<br>L3_logB |
| N703 2 | D<br>L1_gA | D<br>L1_SA | D<br>L1_LB | D<br>L2_4xA | D<br>L2_logB | D<br>L3_gA | D<br>L3_LA | D<br>C2 |
| N704 3 | D<br>L1_invgA | D<br>L1_gB | D<br>L1_MB | D<br>C03 | D<br>L2_fiboB | D<br>L3_invgA | D<br>L3_MA | D<br>L3_fiboB |
| N705 4 | D<br>L1_equA | D<br>L1_invgB | D<br>L1_SB | D<br>L2_XLA | D<br>L2_4xB | D<br>C4 | D<br>L3_SA | D<br>L3_4xB |
| N711 5 | D<br>L1_logA | D<br>C05 | D<br>L2_gA | D<br>L2_LA | D<br>L2_XLB | D<br>L3_equA | D<br>L3_gB | D<br>L3_XLB |
| N712 6 | D<br>L1_fiboA | D<br>L1_equB | D<br>L2_invgA | D<br>L2_MA | D<br>C6 | D<br>L3_logA | D<br>L3_invgB | D<br>L3_LB |
| N714 7 | D<br>L1_4xA | D<br>L1_logB | D<br>C07 | D<br>L2_SA | D<br>L2_LB | D<br>L3_fiboA | D<br>L3_equB | D<br>L3_MB |
| N715 8 | D<br>L1_XLA | D<br>L1_fiboB | D<br>L2_equA | D<br>L2_gB | D<br>L2_MB | D<br>L3_4xA | D<br>C8 | D<br>L3_SB |
| N718 9 | D<br>L1_LA | D<br>L1_4xB | D<br>L2_logA | D<br>L2_invgB | D<br>C9 | C<br>101<br>101_3 | C<br>Kit_M<br>Kit_M3 | D<br>118abd<br>Abd_mock4 |
| N723 10 | A<br>A_M<br>A_M1 | A<br>103<br>103_1 | A<br>Kit_M<br>Kit_M1 | B<br>103<br>103_2 | B<br>44.1<br>GBOL_mock2 | C<br>103<br>103_3 | D<br>A_M<br>A_M4 | C<br>C10 |
| N724 11 | A<br>C11 | A<br>118abd<br>Abd_mock1 | B<br>A_M<br>A_M2 | B<br>118abd<br>Abd_mock2 | B<br>Kit_M<br>Kit_M2 | C<br>118abd<br>Abd_mock3 | D<br>101<br>101_4 | D<br>44.1<br>GBOL_mock4 |
| N727 12 | A<br>101<br>101_1 | A<br>44.1<br>GBOL_mock1 | B<br>101<br>101_2 | B<br>C12 | C<br>A_M<br>A_M3 | C<br>44.1<br>GBOL_mock3 | D<br>103<br>103_4 | D<br>Kit_M<br>Kit_M4 |

**Fig. S2:** Fig. S2: Plate map for the DNA extraction from the 69 lysate samples (including 9 negative controls), as well as additional wells for metabarcoding primer testing as part of a different project. The letters A-D indicate the primers used in PCR1: **A** BF3 + BR2, **B** mlCOIintF + Fol-degen-rev, **C** mlCOIintF + HCO2198-JJ2, **D** fwhF2 + Fol-de- gen-rev. Additionally, the Illumina indexing used for PCR2 is indicated for each row and column.
