## Supplementary material for "Pooling size sorted malaise trap fractions to maximise taxon recovery with metabarcoding": Figure S3

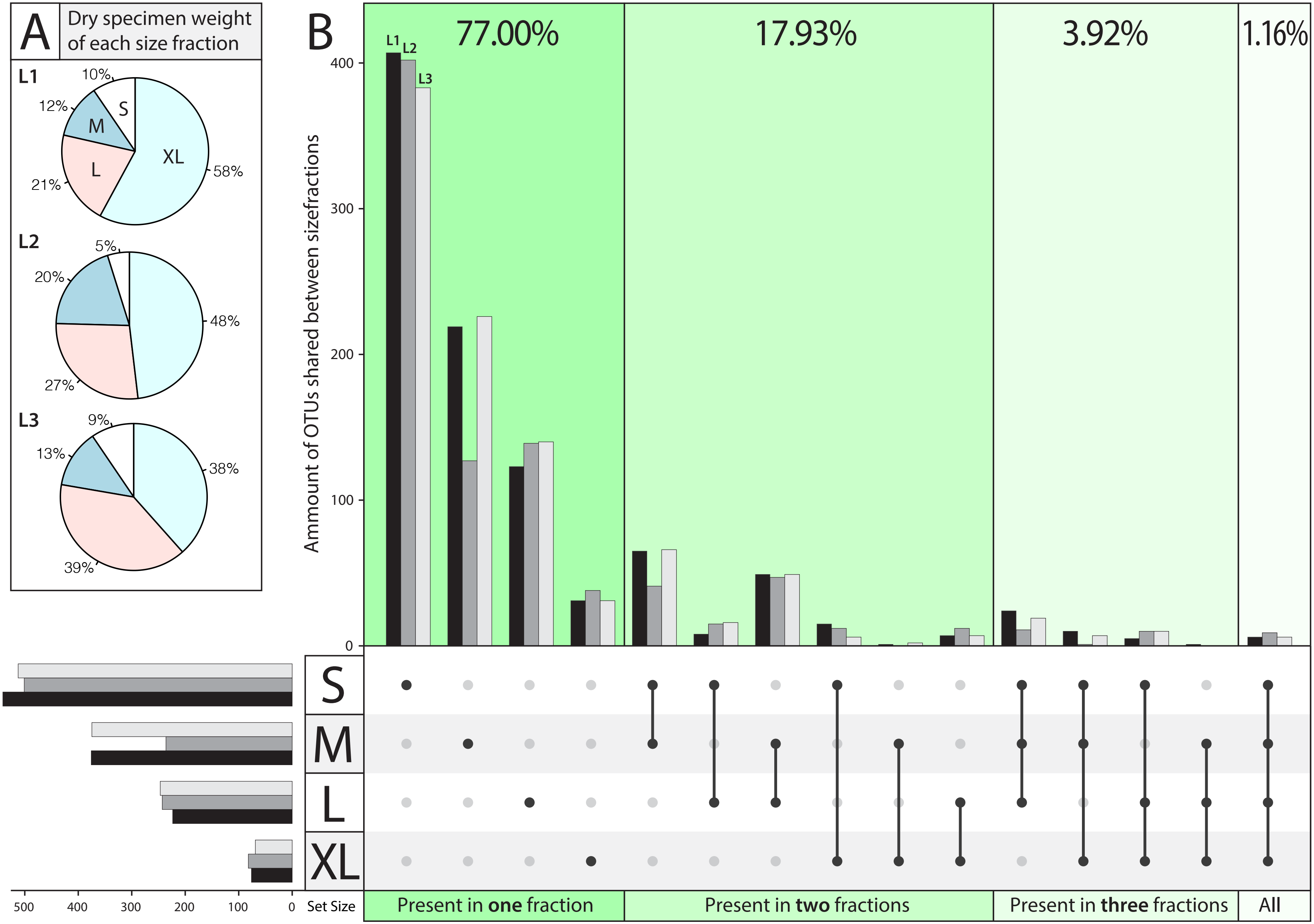


**Fig. S3:** Overview of size fraction dry weight and OTU sharing between size fractions for all 3 malaise trap samples. **A** Pie charts showing the dry specimen weight for each of the 4 individually sequenced size fractions. **B** UpSet plot (Lex 2014) showing the amount of OTUs shared across the 4 size fractions for the 3 samples (L1, L2 and L3, in different shading).
