## Supplementary material for "Pooling size sorted malaise trap fractions to maximise taxon recovery with metabarcoding": Figure S4

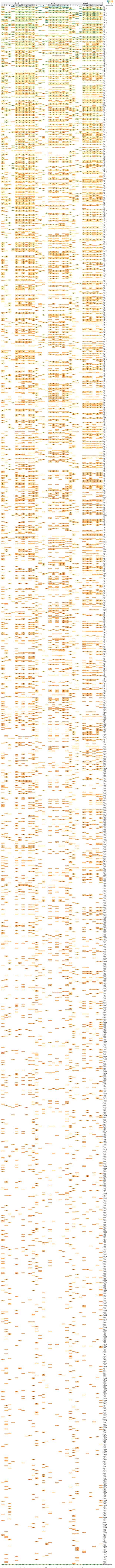

**Fig. S4:** Heatmap indicating the relative read abundance for the different lysates. Replicates were merged, and reads only present in one sample discarded. Reads are subsampled to 10.000 reads per sample (reads with abundance below 0.01% were discarded). See Table S2 for the raw data.
