## Supplementary material for "Pooling size sorted malaise trap fractions to maximise taxon recovery with metabarcoding": Figure S5

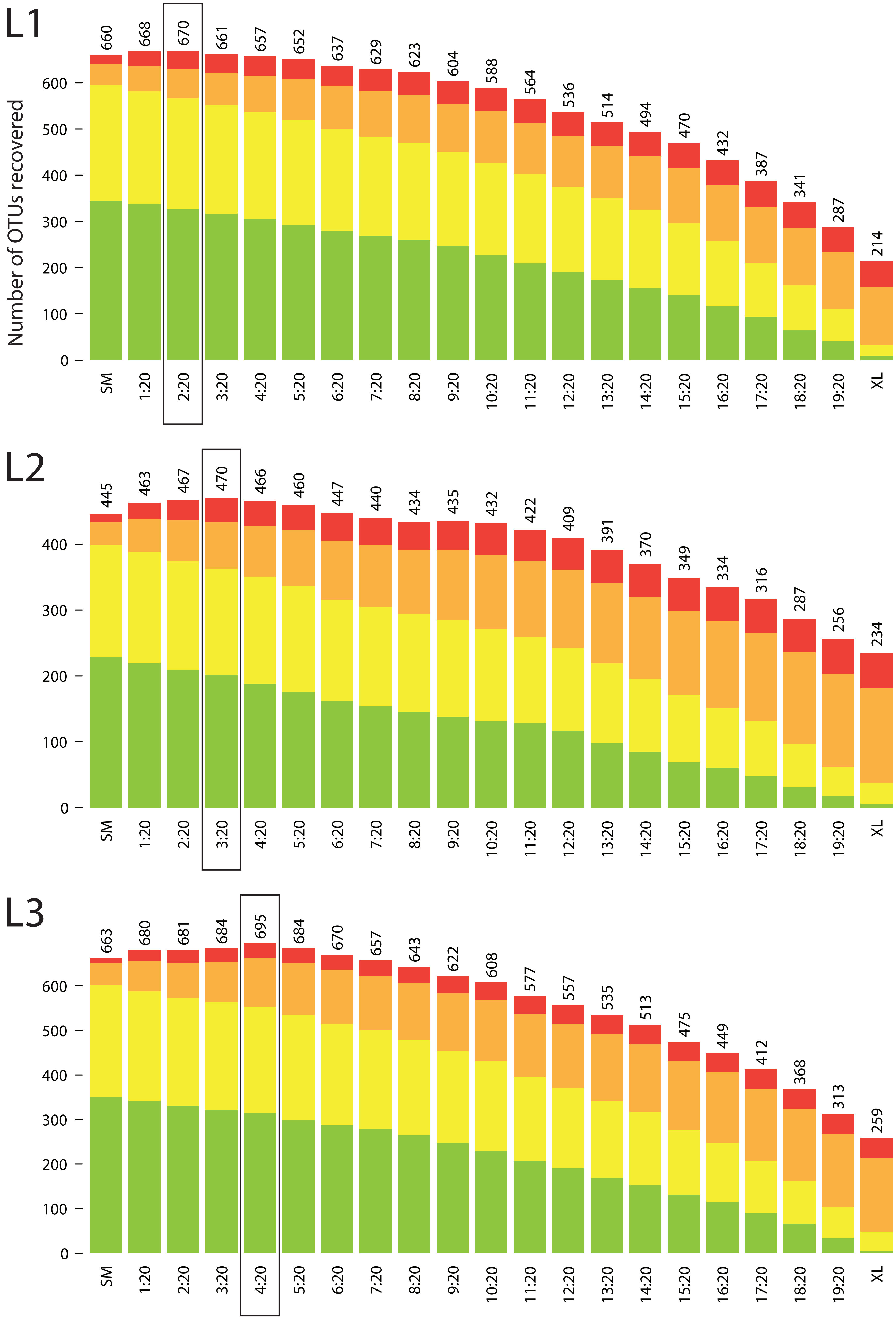


**Fig. S5:** Barplot of OTU recovery for each sample (L1, L2, L3) in silico combined individually sequenced size fraction S+M and L+XL, and the small and large fraction pooled in different proportions. Comparison was done at 10.000 reads sequencing depth.
