## Supplementary material for "Pooling size sorted malaise trap fractions to maximise taxon recovery with metabarcoding": Scripts S1: JAMP_lysis_test_v3.R

```
# 200331 - JAMP processing pimer tests

# install illumina utils
# pip3 install illumina-utils


# trying https://github.com/grenaud/deML
# brew install cmake

deML -i deML.tsv -f Undetermined_S0_L001_R1_001.fastq.gz -r Undetermined_S0_L001_R2_001.fastq.gz -if1 Undetermined_S0_L001_I1_001.fastq.gz  -if2 Undetermined_S0_L001_I2_001.fastq.gz -o test/ --mm 0


#rename files -> fastq

setwd("~/Desktop/primer_test/demulti")


files <- list.files()


file.rename(files, sub("fq$", "fastq", files))


# install jamp 

install.packages(c("bold", "XML", "seqinr", "devtools", "fastqcr"), dependencies=T)
library("devtools")
install_github("VascoElbrecht/PrimerMiner", subdir="PrimerMiner")
install_github("VascoElbrecht/JAMP", subdir="JAMP")


library("JAMP")


# check tiles

setwd("~/Desktop/tiles")


PlotTiles("_101_1_R1_R2.fastq", folder="tile3")


library("JAMP")

setwd("~/Desktop/primer_test/JAMPv2")

# move relevant files for the lysis test over into folder

Empty_folder()

Merge_PE(exe="vsearch", LDist=T)


Cutadapt(forward="GGDACWGGWTGAACWGTWTAYCCHCC", reverse= "TANACYTCNGGRTGNCCRAARAAYCA", LDist=T, error=0.3, cores=12)


Minmax(min=303, max=323, LDist=T)

Max_ee()


# rename files (remove _)
files <- list.files("E_U_max_ee/_data", full.names=T)

file.rename(files, sub("_data/_", "_data/", files))


# clustering only
ee <- list.files("E_U_max_ee/_data", full.names=T)


Cluster_otus(files=ee)

Cluster_otus(files=ee, maxrejects=256, maxaccepts=16)


# getting taxonomy of raw OTU from gbif https://www.gbif.org/tools/sequence-id

# lulu curration of raw data

setwd("~/Documents/ZFMK/2020 lysis + primer test/JAMP")

LULU("OTU_0.01_16_256/3_Raw_OTU_table.csv", folder="200403_lulu_default")


# merge raw data toether

data <- read.csv("OTU_0.01_16_256/1 Raw_lulu_flags.csv", stringsAsFactors=F)

gbif <- read.csv("OTU_0.01_16_256/gbif_200403.csv", stringsAsFactors=F)

head(data)
head(gbif[,c(3,7,8)])

table(data$ID == sub("(.*);size=.*", "\\1", gbif$occurrenceId))


data <- data.frame(data, gbif[,c(3,7,8)], stringsAsFactors=F)


#data <- data[data$parent_id =="OTU_2",]


write.csv(data, "OTU_0.01_16_256/200403_merged_RAW_data.csv", row.names=F)

library("JAMP")
row.names(data) <- data$ID

write.csv(data, "data_cleanup/A 200403_merged_RAW_data.csv", row.names=F)
pdf("data_cleanup/A 200403_merged_RAW_data.pdf", height=(nrow(data)+20)/10, width=(ncol(data)-1)/3)
OTU_heatmap(data[3:71], abundance=T, rel=F, col=rev(c("#d7191c", "#fdae61", "#ffffbf", "#abdda4", "#2b83ba")))
dev.off()


# collapse OTUs based on LULU clasification and assigned taxonomy
# If taxa 

# use for abundance based subsetting!!
rawSums <- colSums(data[3:71])


merged <- data[1,]
merged[1,] <- NA


uniOTU <- unique(data$parent_id)

for(i in 1:length(uniOTU)){
temp <- data[data$parent_id==uniOTU[i],]
temp <- temp[order(rowSums(temp[3:71]), decreasing=T),]

message(length(uniOTU), "/", i)

species <- unique(temp$classification)
for(j in 1:length(species)){
speciestemp <- temp[temp$classification==species[j],]

merged <- rbind(merged, c(speciestemp[1, 1:2], colSums(speciestemp[3:71]), speciestemp[1, 72:81]))
}
}

merged <- merged[-1,]
merged <- merged[order(rowSums(merged[3:71]), decreasing=T),]

row.names(merged) <- merged$ID

write.csv(merged, "data_cleanup/B lulu collapsing.csv", row.names=F)
pdf("data_cleanup/B  lulu collapsing.pdf", height=(nrow(merged)+20)/10, width=(ncol(merged)-1)/3)
OTU_heatmap(merged[3:71], abundance=T, rel=F, col=rev(c("#d7191c", "#fdae61", "#ffffbf", "#abdda4", "#2b83ba")))
dev.off()


sum(merged[3:71])
sum(data[3:71])


# Substract max value *2 of each negative controll from all reads
merged2 <- merged

neg <- merged2[3:11]

sub <- rep(0, nrow(neg))
for(i in 1:nrow(neg)){
sub[i] <- max(neg[i,])*2
}


for(i in 3:71){
merged2[,i] <- merged2[,i] - sub
merged2[,i][merged2[,i]<0] <- 0
}


write.csv(merged2, "data_cleanup/C substr negatives.csv", row.names=F)
pdf("data_cleanup/C substr negatives.pdf", height=(nrow(merged2)+20)/10, width=(ncol(merged2)-1)/3)
OTU_heatmap(merged2[3:71], abundance=T, rel=F, col=rev(c("#d7191c", "#fdae61", "#ffffbf", "#abdda4", "#2b83ba")))
dev.off()


# remove read values below 0.01% abundance
merged3 <- merged2

for(i in 3:71){
merged3[,i] <- round(merged3[,i]/rawSums[i-2]*100, digits=5)
merged2[,i][merged3[,i]<0.01] <- 0
}


write.csv(merged2, "data_cleanup/D 0.01.csv", row.names=F)
pdf("data_cleanup/D 0.01.pdf", height=(nrow(merged3)+20)/10, width=(ncol(merged3)-1)/3)
OTU_heatmap(merged3[3:71], abundance=T, rel=F, col=rev(c("#d7191c", "#fdae61", "#ffffbf", "#abdda4", "#2b83ba")))
dev.off()


# Remove reads that are not present in both replicates
for (i in seq(12, 71, 2)){
A <- merged2[,i]>0
B <- merged2[,i+1]>0
rm <- A+B !=2
merged2[rm,i] <- 0
merged2[rm,i+1] <- 0
}


# remove empty OTUs
merged4 <- merged2[rowSums(merged2[3:71])>0,]

# add discarded read counts to table
discarded <- c(NA, NA, rawSums - colSums(merged4[3:71]), rep(NA, 10) )

merged5 <- rbind(merged4, discarded)
merged5[nrow(merged5), 1:2] <- c(nrow(merged4)+1, "Below_0.01")


write.csv(merged5, "data_cleanup/E replicates checked.csv", row.names=F)
pdf("data_cleanup/E replicates checked.pdf", height=(nrow(merged5)+20)/10, width=(ncol(merged5)-1)/3)
OTU_heatmap(merged5[3:71], abundance=T, rel=F, col=rev(c("#d7191c", "#fdae61", "#ffffbf", "#abdda4", "#2b83ba")))
dev.off()


nrow(merged5)
nrow(merged3)


# pool replicates!

cbind(names(merged5))

merged6 <- merged5[,1:2]
for (i in seq(12, 71, 2)){

merged6 <- data.frame(merged6, merged5[,i]+merged5[, i+1], stringsAsFactors=F)
names(merged6)[ncol(merged6)] <- sub("A$", "", names(merged5[i]))
}


cbind(names(merged6))
# sort
merged6 <- merged6[,c(1,2, 6,5,4,7,3,8:12, c(6,5,4,7,3,8:12)+10, c(6,5,4,7,3,8:12)+20)]


merged6 <- data.frame(merged6, merged5[,72:81], stringsAsFactors=F)

sum(merged5[,12:71])
sum(merged6[,3:32])

# build gbif taxonomy


sub("^(.*)_.*", "\\1", merged6$classification)


A <- strsplit(merged6$classification, "_")
for(i in 1:length(A)){
if(length(A[[i]])<7){A[[i]] <- c(A[[i]], rep(NA, 7-length(A[[i]])))}
}

B <- data.frame(A, stringsAsFactors=F)
names(B) <- 1:ncol(B)
B <- data.frame(t(B), stringsAsFactors=F)

names(B) <- c("Kingdom", "Phylum", "Class", "Order", "Family", "Genus", "Species")

merged6 <- data.frame(merged6, B, stringsAsFactors=F)


# write final tables for stats
write.csv(merged6, "data_cleanup/1 OTU lysis clean.csv", row.names=F)

pdf("data_cleanup/1 OTU lysis clean.pdf", height=(nrow(merged6)+20)/10, width=(ncol(merged6)-1)/3)
OTU_heatmap(merged6[3:32], abundance=T, rel=F, col=rev(c("#d7191c", "#fdae61", "#ffffbf", "#abdda4", "#2b83ba")))
dev.off()

# convert to rel abundnace
merged6rel <- merged6

for(i in 3:32){
merged6rel[,i] <- round(merged6[,i]/sum(merged6[i])*100, digits=5)
merged6rel[,i][merged6rel[,i]<0.01] <- 0
}

head(merged6rel)

sum(merged6[3:32])
sum(data[3:71])

write.csv(merged6rel, "data_cleanup/1 OTU lysis clean rel.csv", row.names=F)

pdf("data_cleanup/1 OTU lysis clean rel.pdf", height=(nrow(merged6rel)+20)/10, width=(ncol(merged6rel)-1)/3)
OTU_heatmap(merged6rel[3:32], abundance=T, rel=F, col=rev(c("#d7191c", "#fdae61", "#ffffbf", "#abdda4", "#2b83ba")))
dev.off()


cat(file="data_cleanup/1 OTU lysis clean.fasta", paste(paste(">", merged6$ID[-nrow(merged6)], sep=""), toupper(merged6$sequ[-nrow(merged6)]), sep="\n"), sep="\n")
```
