## Supplementary material for "Pooling size sorted malaise trap fractions to maximise taxon recovery with metabarcoding": Scripts S1: indiv_vs_pooled_v1.R

```
# 20200419
setwd("~/Documents/ZFMK/2020 lysis + primer test/R scripts/1 recovery indiv vs pooled")


data <- read.csv("~/Google Drive/WritePapers/2020 lysis poolig/1 supporting info/Tab S1 OTU table/Lysis_OTUs_16_256_v3.csv", stringsAsFactors=F)


# sequencind depth 10.000 = 0.01% abundance, or more
# sequencind depth 2.500 = 0.04% abundance, or more


cbind(names(data))


treshhold <- c(NA, NA, rep(c(0.04, 0.04, 0.04, 0.04, 0.01, 0.01, 0.01, 0.01, 0.01, 0.01), 3) )


for(i in 3:32){
temprel <- data[,i]/sum(data[,i])*100 
data[,i] <- temprel
data[,i][temprel < treshhold[i]] <- 0

data[length(temprel), i] <- 100 - sum(temprel[-length(temprel)][temprel[-length(temprel)] >= treshhold[i]])
}


colSums(data[3:32])
tail(data[,3:32])


L1 <- rowSums(data[3:6])
L1 <- L1/sum(L1)*100


L2 <- rowSums(data[13:16])
L2 <- L2/sum(L2)*100


L3 <- rowSums(data[23:26])
L3 <- L3/sum(L3)*100


temp <- data.frame(L1, L2, L3, "size_estimate"=data$size_estimate, stringsAsFactors=F)


# plot OTUs recovered
data2 <- temp[-nrow(temp),]

size_estimate <- data2$size_estimate
size_estimate[is.na(size_estimate)] <- "na"


# absolute read abundance
temp <- data.frame("Sample"=NA, "S"=NA, "M"=NA, "L"=NA, "XL"=NA, "na"=NA, stringsAsFactors=F)

for (i in 1:3){
temp[i,1] <- names(data2)[i]
temp[i,2] <- sum(data2[,i][size_estimate=="S"]>0)
temp[i,3] <- sum(data2[,i][size_estimate=="M"]>0)
temp[i,4] <- sum(data2[,i][size_estimate=="L"]>0)
temp[i,5] <- sum(data2[,i][size_estimate=="XL"]>0)
temp[i,6] <- sum(data2[,i][size_estimate=="na"]>0)

#temp[i, 2:6] <- temp[i, 2:6]/sum(temp[i, 2:6])*100 # convert to rel abundance
}

#temp <- temp[-1,]


# make plot with absolute abundance
pdf("OTU_indivpooled_14.pdf", width=4, height=6)
x <- barplot(as.matrix(t(temp[,-1])), xaxt="n", yaxt="n", border=NA, col=c("#8DC63F", "#F9ED32", "#FBB040", "#EF4136", "#BCBEC0"), ylim=c(0,600))
axis(2, las=1)
#axis(1, las=2, at=x, sub("L._", "", temp$Sample))
axis(1, las=2, at=x, temp$Sample, tick=F)
text(x, colSums(as.matrix(t(temp[,-1])))+10, colSums(as.matrix(t(temp[,-1]))), srt=90, adj=0)
dev.off()

write.csv(temp, "OTU_recovery_stats.csv")


# relative proportions


L1 <- rowSums(data[3:6])
L1 <- L1/sum(L1)*100


L2 <- rowSums(data[13:16])
L2 <- L2/sum(L2)*100


L3 <- rowSums(data[23:26])
L3 <- L3/sum(L3)*100


temp <- data.frame(L1, L2, L3, "size_estimate"=data$size_estimate, stringsAsFactors=F)


data <- temp[-nrow(temp),]

size_estimate <- data$size_estimate
size_estimate[is.na(size_estimate)] <- "na"


# absolute read abundance
temp <- data.frame("Sample"=NA, "S"=NA, "M"=NA, "L"=NA, "XL"=NA, "na"=NA, stringsAsFactors=F)

for (i in 1:3){
temp[i,1] <- names(data)[i]
temp[i,2] <- sum(data[,i][size_estimate=="S"])
temp[i,3] <- sum(data[,i][size_estimate=="M"])
temp[i,4] <- sum(data[,i][size_estimate=="L"])
temp[i,5] <- sum(data[,i][size_estimate=="XL"])
temp[i,6] <- sum(data[,i][size_estimate=="na"])

temp[i, 2:6] <- temp[i, 2:6]/sum(temp[i, 2:6])*100 # convert to rel abundance
}

#temp <- temp[-1,]


pdf("OTU_indivpooled_14_rel.pdf", width=4, height=6)
x <- barplot(as.matrix(t(temp[,-1])), xaxt="n", yaxt="n", border=NA, col=c("#8DC63F", "#F9ED32", "#FBB040", "#EF4136", "#BCBEC0"), ylim=c(0,100))
axis(2, las=1)
#axis(1, las=2, at=x, sub("L._", "", temp$Sample))
axis(1, las=2, at=x, temp$Sample, tick=F)
dev.off()
```
