## Supplementary material for "Pooling size sorted malaise trap fractions to maximise taxon recovery with metabarcoding": Scripts S1: lysis_first_look_v1.R

```
# 200403 first look at lysis data

setwd("~/Documents/ZFMK/2020 lysis + primer test/R scripts/1 first look")

data <- read.csv("~/Documents/ZFMK/2020 lysis + primer test/JAMP/data_cleanup/1 OTU lysis clean.csv", stringsAsFactors=F)

nrow(data)

data <- data[data$identity>=98,]
data <- data[!is.na(data$identity),]

nrow(data)


cbind(names(data))


str(data[,12:71])


pdf("OTU_counts.pdf", width=14, height=10)
x <- barplot(colSums(data[,12:71]>0), col=c("Gray"), xaxt='n', ylim=c(0, 780))

axis(1, at=x, names(data[,12:71]), las=2)
text(x,colSums(data[,12:71]>0)+20, colSums(data[,12:71]>0)+10, srt=90)


dev.off()


# check replicates after 00.0% filtering
rep <- read.csv("~/Documents/ZFMK/2020 lysis + primer test/JAMP/data_cleanup/D 0.01.csv", stringsAsFactors=F)

pdf("replication.pdf")
for(i in seq(12, 71, 2)){
plot(rep[,i]/sum(rep[,i])*100, rep[,i+1]/sum(rep[,i+1])*100, log="xy", main=sub("[AB]$", "", names(rep)[i]), xlab=names(rep)[i], ylab=names(rep)[i+1], xaxt="n", yaxt="n", xlim=c(0.01, 100), ylim=c(0.01, 100))
axis(1, at=c(0.01, 0.1, 1, 10, 100), labels=c("0.01", "0.1", "1", "10", "100"))
axis(2, at=c(0.01, 0.1, 1, 10, 100), labels=c("0.01", "0.1", "1", "10", "100"))

lines(c(0.005, 100), c(0.005, 100), col="Red")

}
dev.off()
```
