## Supplementary material for "Pooling size sorted malaise trap fractions to maximise taxon recovery with metabarcoding": Scripts S1: rarefaction_v1.R

```
# 200405 rarefaction

setwd("~/Documents/ZFMK/2020 lysis + primer test/R scripts/1 rarefaction")

# processed table - limited to 0.01%, pick neg control filtered table instead.
#data <- read.csv("~/Google Drive/WritePapers/2020 lysis poolig/1 supporting info/Tab S1 OTU table/Lysis_OTUs_16_256_v3_rel.csv", stringsAsFactors=F)


data <- read.csv("~/Documents/ZFMK/2020 lysis + primer test/JAMP/data_cleanup/C substr negatives.csv", stringsAsFactors=F)


cbind(names(data))
row.names(data) <- data$ID
data <- data[, c(1:2, 12:71)]


data2 <- data
# pool replicates! and convert to rel abundance

data2 <- data[,1:2]

cbind(names(data))

for (i in seq(3, 62, 2)){
# merge
temp <- data[,i]+ data[, i+1]
# rel abundance
temp <- round(temp/sum(temp)*100, 6)

# add to data frame
data2 <- data.frame(data2, temp, stringsAsFactors=F)
names(data2)[ncol(data2)] <- sub("A$", "", names(data[i]))
}


nrow(data2)


data <- data2[,-c(1,2)]


tail(data)
head(data)


# 0.01% = 10.000 sequences
# 0.02% = 5.000 sequences
# 0.1% = 1.000 sequences

depth <- c(rev(seq(10000, 50000, by=10000)), rev(seq(1000, 10000, by=1000)), rev(seq(500, 900, by=100)))


all <- NULL

for(i in 100/depth){
exp <- NULL

for (j in 1:ncol(data)){
exp[j] <- sum(data[,j]> i)
}

all <- cbind(all, exp)
}


temp <- data.frame("sample"=names(data), all, stringsAsFactors=F)

names(temp) <- c("sample", paste("V", depth, sep=""))


write.csv(temp, "sequdepth.csv")


temp <- temp[-c(2:5, 12:15, 22:25), ]

col <- rep(c("Red", "Darkgray", "Green", "Black", "Blue", "Orange"), 3)


temp3 <- temp[c(1:6), ]
pdf("L1.pdf", height=9, width=6)
plot(NULL, log="x", ylim=c(0, 1600), xlim=c(500, 70000))

for (i in 1:nrow(temp3)){
lines(depth, temp3[i,-1], col=col[i], lwd=2)
text(55000, temp3[i,2], sub("L._", "", temp3$sample[i]), adj=0, cex=1, col=col[i])
}
dev.off()


temp3 <- temp[c(7:12), ]
pdf("L2.pdf", height=9, width=6)
plot(NULL, log="x", ylim=c(0, 1600), xlim=c(500, 70000))

for (i in 1:nrow(temp3)){
lines(depth, temp3[i,-1], col=col[i], lwd=2)
text(55000, temp3[i,2], sub("L._", "", temp3$sample[i]), adj=0, cex=1, col=col[i])
}
dev.off()


temp3 <- temp[c(13:18), ]
pdf("L3.pdf", height=9, width=6)
plot(NULL, log="x", ylim=c(0, 1600), xlim=c(500, 70000))

for (i in 1:nrow(temp3)){
lines(depth, temp3[i,-1], col=col[i], lwd=2)
text(55000, temp3[i,2], sub("L._", "", temp3$sample[i]), adj=0, cex=1, col=col[i])
}
dev.off()
```
