## Supplementary material for "Pooling size sorted malaise trap fractions to maximise taxon recovery with metabarcoding": Scripts S1: silico_pool_v1.R

```
#20200421

setwd("~/Documents/ZFMK/2020 lysis + primer test/R scripts/1 in silico pool")


# Add insect weight
# S, M, L, XL
L1 <- c(1.1832, 1.4836, 2.5662, 7.2005)
L2 <- c(0.4772, 1.9375, 2.6814, 4.7291)
L3 <- c(1.0076, 1.3586, 4.1691, 4.0783)

L1 <- L1/sum(L1)*100
L2 <- L2/sum(L2)*100
L3 <- L3/sum(L3)*100


data <- read.csv("~/Google Drive/WritePapers/2020 lysis poolig/1 supporting info/Tab S1 OTU table/Lysis_OTUs_16_256_v3_rel.csv", stringsAsFactors=F)

# pool S and L based on weight
L1s <- L1[1:2]/sum(L1[1:2])

L1sm <- data$L1_S*L1s[1] + data$L1_M*L1s[2]
L1sm <- L1sm/sum(L1sm)*100

#cbind(L1sm, data$L1_S, data$L1_M)

# pool L and XL based on weight
L1l <- L1[3:4]/sum(L1[3:4])

L1lxl <- data$L1_L*L1l[1] + data$L1_XL* L1l[2]
L1lxl <- L1lxl/sum(L1lxl)*100

#cbind(L1lxl, data$L1_L, data$L1_XL)

# pool L1sm and L1lxl in different proportions

i =11
exp <- NULL
for (i in c(seq(0, 10, 0.5))){
temp <- L1sm*(10-i)+L1lxl*(i)
temp <- temp/sum(temp)*100
temp[temp<0.01] <- 0
#temp <- temp>=0.01
exp <- cbind(exp, temp)
}

exp <- data.frame(exp)

#barplot(colSums(exp>0))

size_estimate <- data$size_estimate
size_estimate[is.na(size_estimate)] <- "na"


exp <- exp[-nrow(exp),]


# absolute read abundance
temp <- data.frame("Sample"=NA, "S"=NA, "M"=NA, "L"=NA, "XL"=NA, "na"=NA, stringsAsFactors=F)

for (i in 1:ncol(exp)){
temp[i,1] <- names(exp)[i]
temp[i,2] <- sum(exp[,i][size_estimate=="S"]>0)
temp[i,3] <- sum(exp[,i][size_estimate=="M"]>0)
temp[i,4] <- sum(exp[,i][size_estimate=="L"]>0)
temp[i,5] <- sum(exp[,i][size_estimate=="XL"]>0)
temp[i,6] <- sum(exp[,i][size_estimate=="na"]>0)

#temp[i, 2:6] <- temp[i, 2:6]/sum(temp[i, 2:6])*100 # convert to rel abundance
}

#temp <- temp[-1,]


if(F){
temp <- data.frame("Sample"=NA, "S"=NA, "M"=NA, "L"=NA, "XL"=NA, "na"=NA, stringsAsFactors=F)

for (i in 1:ncol(exp)){
temp[i,1] <- names(exp)[i]
temp[i,2] <- sum(exp[,i][size_estimate=="S"])
temp[i,3] <- sum(exp[,i][size_estimate=="M"])
temp[i,4] <- sum(exp[,i][size_estimate=="L"])
temp[i,5] <- sum(exp[,i][size_estimate=="XL"])
temp[i,6] <- sum(exp[,i][size_estimate=="na"])

temp[i, 2:6] <- temp[i, 2:6]/sum(temp[i, 2:6])*100 # convert to rel abundance
}

}

temp <- temp[,-6]

text <- c("SM", paste(1:19, ":", 20, sep=""), "XL")

# make plot with absolute abundance
pdf("SM+XL_L1.pdf", width=10, height=6)
x <- barplot(as.matrix(t(temp[,-1])), xaxt="n", yaxt="n", border=NA, col=c("#8DC63F", "#F9ED32", "#FBB040", "#EF4136", "#BCBEC0"))
axis(2, las=1)
#axis(1, las=2, at=x, sub("L._", "", temp$Sample))
axis(1, las=2, at=x, text, tick=F)

text(x, colSums(as.matrix(t(temp[,-1])))+10, colSums(as.matrix(t(temp[,-1]))), srt=90, adj=0)

dev.off()


# pool S and L based on weight
L2s <- L2[1:2]/sum(L2[1:2])

L2sm <- data$L2_S*L2s[1] + data$L2_M*L2s[2]
L2sm <- L2sm/sum(L2sm)*100

#cbind(L2sm, data$L2_S, data$L2_M)

# pool L and XL based on weight
L2l <- L2[3:4]/sum(L2[3:4])

L2lxl <- data$L2_L*L2l[1] + data$L2_XL* L2l[2]
L2lxl <- L2lxl/sum(L2lxl)*100

#cbind(L2lxl, data$L2_L, data$L2_XL)

# pool L2sm and L2lxl in different proportions

i =11
exp <- NULL
for (i in c(seq(0, 10, 0.5))){
temp <- L2sm*(10-i)+L2lxl*(i)
temp <- temp/sum(temp)*100
temp[temp<0.01] <- 0
#temp <- temp>=0.01
exp <- cbind(exp, temp)
}

exp <- data.frame(exp)

#barplot(colSums(exp>0))

size_estimate <- data$size_estimate
size_estimate[is.na(size_estimate)] <- "na"


exp <- exp[-nrow(exp),]


# absolute read abundance
temp <- data.frame("Sample"=NA, "S"=NA, "M"=NA, "L"=NA, "XL"=NA, "na"=NA, stringsAsFactors=F)

for (i in 1:ncol(exp)){
temp[i,1] <- names(exp)[i]
temp[i,2] <- sum(exp[,i][size_estimate=="S"]>0)
temp[i,3] <- sum(exp[,i][size_estimate=="M"]>0)
temp[i,4] <- sum(exp[,i][size_estimate=="L"]>0)
temp[i,5] <- sum(exp[,i][size_estimate=="XL"]>0)
temp[i,6] <- sum(exp[,i][size_estimate=="na"]>0)

#temp[i, 2:6] <- temp[i, 2:6]/sum(temp[i, 2:6])*100 # convert to rel abundance
}

#temp <- temp[-1,]


temp <- temp[,-6]

text <- c("SM", paste(1:19, ":", 20, sep=""), "XL")

# make plot with absolute abundance
pdf("SM+XL_L2.pdf", width=10, height=6)
x <- barplot(as.matrix(t(temp[,-1])), xaxt="n", yaxt="n", border=NA, col=c("#8DC63F", "#F9ED32", "#FBB040", "#EF4136", "#BCBEC0"))
axis(2, las=1)
#axis(1, las=2, at=x, sub("L._", "", temp$Sample))
axis(1, las=2, at=x, text, tick=F)

text(x, colSums(as.matrix(t(temp[,-1])))+10, colSums(as.matrix(t(temp[,-1]))), srt=90, adj=0)

dev.off()


# pool S and L based on weight
L3s <- L3[1:2]/sum(L3[1:2])

L3sm <- data$L3_S*L3s[1] + data$L3_M*L3s[2]
L3sm <- L3sm/sum(L3sm)*100

#cbind(L3sm, data$L3_S, data$L3_M)

# pool L and XL based on weight
L3l <- L3[3:4]/sum(L3[3:4])

L3lxl <- data$L3_L*L3l[1] + data$L3_XL* L3l[2]
L3lxl <- L3lxl/sum(L3lxl)*100

#cbind(L3lxl, data$L3_L, data$L3_XL)

# pool L3sm and L3lxl in different proportions

i =11
exp <- NULL
for (i in c(seq(0, 10, 0.5))){
temp <- L3sm*(10-i)+L3lxl*(i)
temp <- temp/sum(temp)*100
temp[temp<0.01] <- 0
#temp <- temp>=0.01
exp <- cbind(exp, temp)
}

exp <- data.frame(exp)

#barplot(colSums(exp>0))

size_estimate <- data$size_estimate
size_estimate[is.na(size_estimate)] <- "na"


exp <- exp[-nrow(exp),]


# absolute read abundance
temp <- data.frame("Sample"=NA, "S"=NA, "M"=NA, "L"=NA, "XL"=NA, "na"=NA, stringsAsFactors=F)

for (i in 1:ncol(exp)){
temp[i,1] <- names(exp)[i]
temp[i,2] <- sum(exp[,i][size_estimate=="S"]>0)
temp[i,3] <- sum(exp[,i][size_estimate=="M"]>0)
temp[i,4] <- sum(exp[,i][size_estimate=="L"]>0)
temp[i,5] <- sum(exp[,i][size_estimate=="XL"]>0)
temp[i,6] <- sum(exp[,i][size_estimate=="na"]>0)

#temp[i, 2:6] <- temp[i, 2:6]/sum(temp[i, 2:6])*100 # convert to rel abundance
}

#temp <- temp[-1,]


temp <- temp[,-6]

text <- c("SM", paste(1:19, ":", 20, sep=""), "XL")

# make plot with absolute abundance
pdf("SM+XL_L3.pdf", width=10, height=6)
x <- barplot(as.matrix(t(temp[,-1])), xaxt="n", yaxt="n", border=NA, col=c("#8DC63F", "#F9ED32", "#FBB040", "#EF4136", "#BCBEC0"))
axis(2, las=1)
#axis(1, las=2, at=x, sub("L._", "", temp$Sample))
axis(1, las=2, at=x, text, tick=F)

text(x, colSums(as.matrix(t(temp[,-1])))+10, colSums(as.matrix(t(temp[,-1]))), srt=90, adj=0)

dev.off()
```
