## Supplementary material for "Pooling size sorted malaise trap fractions to maximise taxon recovery with metabarcoding": Scripts S1: size_assign_v3.R

```
# 200404 - assign size estimate S, M, L, XL to OTUs based on individualy seqeucned samples.

setwd("~/Documents/ZFMK/2020 lysis + primer test/R scripts/1 assign size")

data <- read.csv("~/Documents/ZFMK/2020 lysis + primer test/JAMP/data_cleanup/1 OTU lysis clean rel.csv", stringsAsFactors=F)

head(data)

A <- data[,3:6]
B <- data[,13:16]
C <- data[,23:26]

temp <- A+B+C

names(temp) <- c("S", "M", "L", "XL")
row.names(temp) <- data$ID

temp <- temp[-nrow(temp),]


size_estimate <- NULL
for (i in 1:nrow(temp)){
if(sum(temp[i,])>0){
size_estimate[i] <- names(temp)[max(temp[i,])==temp[i,]]
} else {
size_estimate[i] <- NA
}
}


temp2 <- temp
row.names(temp2) <- paste(size_estimate, "  ", row.names(temp2))


library("JAMP")

pdf("combined_relSMLXL.pdf", height=(nrow(temp)+20)/10, width=(ncol(temp)+4)/3)
OTU_heatmap(round(temp2,2), abundance=T, rel=F, col=rev(c("#d7191c", "#fdae61", "#ffffbf", "#abdda4", "#2b83ba")))
dev.off()


write.csv(data.frame(temp, size_estimate, stringsAsFactors=F), file="SMLXL.csv")


# 200404 - make plots of taxa detection based on size charts

data <- read.csv("~/Google Drive/WritePapers/2020 lysis poolig/1 supporting info/Tab S1 OTU table/Lysis_OTUs_16_256_v3.csv", stringsAsFactors=F)

nrow(data)
data <- data[-nrow(data),]
nrow(data)

cbind(names(data))
size_estimate <- data$size_estimate
size_estimate[is.na(size_estimate)] <- "na"

data <- data[,c(2:32, 50)]
head(data)


# absolute read abundance
temp <- data.frame("Sample"=NA, "S"=NA, "M"=NA, "L"=NA, "XL"=NA, "na"=NA, stringsAsFactors=F)

for (i in 2:31){
temp[i,1] <- names(data)[i]
temp[i,2] <- sum(data[,i][size_estimate=="S"])
temp[i,3] <- sum(data[,i][size_estimate=="M"])
temp[i,4] <- sum(data[,i][size_estimate=="L"])
temp[i,5] <- sum(data[,i][size_estimate=="XL"])
temp[i,6] <- sum(data[,i][size_estimate=="na"])

temp[i, 2:6] <- temp[i, 2:6]/sum(temp[i, 2:6])*100 # convert to rel abundance
}

temp <- temp[-1,]


cbind(temp$Sample)

#temp <- temp[c(1,11,21, 2,12,22, 3,13,23, 4,14,24, 8,18,28, 6,16,26, 9,19,29, 7,17,27, 5,15,25, 10,20,30),]
temp <- temp[c(1,11,21, 2,12,22, 3,13,23, 4,14,24, 8,6,9,7,5,10, 18,16,19,17,15,20, 28,26,29,27,25,30),]


# Add insect weight
L1 <- c(1.1832, 1.4836, 2.5662, 7.2005)
L2 <- c(0.4772, 1.9375, 2.6814, 4.7291)
L3 <- c(1.0076, 1.3586, 4.1691, 4.0783)

L1 <- L1/sum(L1)*100
L2 <- L2/sum(L2)*100
L3 <- L3/sum(L3)*100


dry <- data.frame(c("L1_dry", "L2_dry", "L3_dry"), rbind(L1, L2, L3), 0)
names(dry) <- names(temp)

temp2 <- rbind(temp[1:12,], dry, temp[13:30,])


write.csv(temp2, "recovery_stats_AbsAbundance_20200419.csv")


# make plot with absolute abundance
pdf("Abs_taxa_recovery(readcounts)+20200419.pdf", width=10, height=6)
x <- barplot(as.matrix(t(temp2[,-1])), xaxt="n", yaxt="n", border=NA, col=c("#8DC63F", "#F9ED32", "#FBB040", "#EF4136", "#BCBEC0"))
axis(2, las=1)
#axis(1, las=2, at=x, sub("L._", "", temp$Sample))
axis(1, las=2, at=x, temp2$Sample, tick=F)
dev.off()


# plot OTUs recovered
data2 <- data[-nrow(data),]

size_estimate <- data2$size_estimate
size_estimate[is.na(size_estimate)] <- "na"


# absolute read abundance
temp <- data.frame("Sample"=NA, "S"=NA, "M"=NA, "L"=NA, "XL"=NA, "na"=NA, stringsAsFactors=F)

for (i in 2:31){
temp[i,1] <- names(data2)[i]
temp[i,2] <- sum(data2[,i][size_estimate=="S"]>0)
temp[i,3] <- sum(data2[,i][size_estimate=="M"]>0)
temp[i,4] <- sum(data2[,i][size_estimate=="L"]>0)
temp[i,5] <- sum(data2[,i][size_estimate=="XL"]>0)
temp[i,6] <- sum(data2[,i][size_estimate=="na"]>0)

#temp[i, 2:6] <- temp[i, 2:6]/sum(temp[i, 2:6])*100 # convert to rel abundance
}

temp <- temp[-1,]


cbind(temp$Sample)

#temp <- temp[c(1,11,21, 2,12,22, 3,13,23, 4,14,24, 8,18,28, 6,16,26, 9,19,29, 7,17,27, 5,15,25, 10,20,30),]
temp <- temp[c(1,11,21, 2,12,22, 3,13,23, 4,14,24, 8,6,9,7,5,10, 18,16,19,17,15,20, 28,26,29,27,25,30),]


# Add max detected number of OTUs 

L1 <- size_estimate[rowSums(data2[2:11])>0]
L1 <- table(L1)
L1 <- c(L1[4], L1[2], L1[1], L1[5], L1[3])


L2 <- size_estimate[rowSums(data2[12:21])>0]
L2 <- table(L2)
L2 <- c(L2[4], L2[2], L2[1], L2[5], L2[3])

L3 <- size_estimate[rowSums(data2[22:31])>0]
L3 <- table(L3)
L3 <- c(L3[4], L3[2], L3[1], L3[5], L3[3])


# deactivate
L1 <- c(0,0,0,0,0)
L2 <- c(0,0,0,0,0)
L3 <- c(0,0,0,0,0)


dry <- data.frame(c("L1_dry", "L2_dry", "L3_dry"), rbind(L1, L2, L3))
names(dry) <- names(temp)

temp2 <- rbind(temp[1:12,], dry, temp[13:30,])


# make plot with absolute abundance
pdf("OTU_recovery+.pdf", width=10, height=6)
x <- barplot(as.matrix(t(temp2[,-1])), xaxt="n", yaxt="n", border=NA, col=c("#8DC63F", "#F9ED32", "#FBB040", "#EF4136", "#BCBEC0"), ylim=c(0,600))
axis(2, las=1)
#axis(1, las=2, at=x, sub("L._", "", temp$Sample))
axis(1, las=2, at=x, temp2$Sample, tick=F)
text(x, colSums(as.matrix(t(temp2[,-1])))+10, colSums(as.matrix(t(temp2[,-1]))), srt=90, adj=0)
dev.off()

write.csv(temp2, "OTU_recovery_stats.csv")


# 20200419

S <- colSums(temp2[1:3,-1])
M <- colSums(temp2[4:6,-1])
L <- colSums(temp2[7:9,-1])
XL <- colSums(temp2[10:12,-1])


average <- c(sum(S)/3, sum(M)/3, sum(L)/3, sum(XL)/3)


round(average/sum(average)*100, 2)


# count specimen sizes

data <- read.csv("~/Google Drive/WritePapers/2020 lysis poolig/1 supporting info/Tab S1 OTU table/Lysis_OTUs_16_256_v3.csv", stringsAsFactors=F)

mysize <- data$size_estimate[-nrow(data)]
mysize[is.na(mysize)] <- "NA"


mysize <- table(mysize[-nrow(data)])

#  L   M  NA   S  XL 
# 422 539 206 912 118 


round(mysize/sum(mysize)*100, 2)


sum <- table(data$Phylum[-nrow(data)], useNA="always")

round(sum/sum(sum)*100)
```
