## Supplementary material for "Pooling size sorted malaise trap fractions to maximise taxon recovery with metabarcoding": Scripts S1: UpSet_v1.R

```
# 200404 UpSet plot - sharing between size fractions


setwd("~/Documents/ZFMK/2020 lysis + primer test/R scripts/1 UpSet plots")

#install.packages("UpSetR", dependencies=T)

library("UpSetR")


data <- read.csv("~/Documents/ZFMK/2020 lysis + primer test/JAMP/data_cleanup/1 OTU lysis clean.csv", stringsAsFactors=F)


A <- data[,3:6]
A <- A>0
A <- A*1
A <- data.frame(A)


cbind(names(data))

B <- data[,13:16]
B <- B>0
B <- B*1
B <- data.frame(B)


C <- data[,23:26]
C <- C>0
C <- C*1
C <- data.frame(C)


pdf("A.pdf", width=10, useDingbats=F)
upset(A, empty.intersections=T, mainbar.y.max=420, keep.order=T, sets=c("L1_S", "L1_M", "L1_L", "L1_XL")[4:1], order.by="degree", decreasing=F)
dev.off()


pdf("B.pdf", width=10)
upset(B, empty.intersections=T, mainbar.y.max=420, keep.order=T, sets=c("L2_S", "L2_M", "L2_L", "L2_XL")[4:1], order.by="degree", decreasing=F)
dev.off()


pdf("C.pdf", width=10)
upset(C, empty.intersections=T, mainbar.y.max=420, keep.order=T, sets=c("L3_S", "L3_M", "L3_L", "L3_XL")[4:1], order.by="degree", decreasing=F)
dev.off()


# calculate percentage in each category
# across all samples = all OTUs!

A <- data[,3:6]
B <- data[,13:16]
C <- data[,23:26]


temp <- A+B+C

names(temp) <- c("V1", "V2", "V3", "V4")
temp <- temp[-nrow(temp),]

temp <- temp>0
temp <- temp*1

temp <- data.frame(temp, stringsAsFactors=F)

temp <- cbind(table(paste(temp$V1, temp$V2, temp$V3, temp$V4, sep="")))


cbind(row.names(temp))

# one fraction only -> 77.00%
round(sum(temp[c(2,3,5,9)])/sum(temp[-1])*100, 2)


# two only -> 17.93%
round(sum(temp[c(4, 6, 7, 10, 11, 13)])/sum(temp[-1])*100, 2)


# 3 only -> 3.92%
round(sum(temp[c(8,12,14,15)])/sum(temp[-1])*100, 2)


# all -> 1.16%
round(sum(temp[c(16)])/sum(temp[-1])*100, 2)


nrow(data)
```
