## Supplementary material for "Pooling size sorted malaise trap fractions to maximise taxon recovery with metabarcoding": Scripts S1: wett_sieve_v2.R

```
# 20200504 - wett sieving size distributions


setwd("~/Documents/ZFMK/2020 lysis + primer test/R scripts/1 size sort")

data <- read.csv("20200507_SL_ÖFS-059-Vlotho.csv", stringsAsFactors=F,  dec=",")


names(data) <- c("Fraction","Order", "body", "antenna", "totallength", "g", "mg")
data$antenna[is.na(data$antenna)] <- 0


head(data)


pdf("specimen_size_small.pdf", height=6)
hist(data$body[data$Fraction=="S"], breaks=0:20, xlim=c(0,20), col="gray", border=F, ylim=c(0, 250))
lines(c(mean(data$body[data$Fraction=="S"]),mean(data$body[data$Fraction=="S"])), c(0, 250))
dev.off()

round(mean(data$body[data$Fraction=="S"]), 2)

length(data$body[data$Fraction=="S"])


pdf("specimen_size_large.pdf", height=6)
hist(data$body[data$Fraction=="L"], breaks=0:20, xlim=c(0,20), col="gray", border=F, ylim=c(0, 250))
lines(c(mean(data$body[data$Fraction=="L"]),mean(data$body[data$Fraction=="L"])), c(0, 250))
dev.off()

round(mean(data$body[data$Fraction=="L"]), 2)


length(data$body[data$Fraction=="L"])


pdf("specimen_weight_small.pdf", height=6)
hist(data$mg[data$Fraction=="S"], breaks=0:70, xlim=c(0,70), col="gray", border=F, ylim=c(0, 500))
lines(c(mean(data$mg[data$Fraction=="S"]),mean(data$mg[data$Fraction=="S"])), c(0, 500))
dev.off()

round(mean(data$mg[data$Fraction=="S"]), 2)


pdf("specimen_weight_large.pdf", height=6)
hist(data$mg[data$Fraction=="L"], breaks=0:70, xlim=c(0,70), col="gray", border=F, ylim=c(0, 500))
lines(c(mean(data$mg[data$Fraction=="L"]),mean(data$mg[data$Fraction=="L"])), c(0, 500))
dev.off()

round(mean(data$mg[data$Fraction=="L"]), 2)

max(data$mg[data$Fraction=="L"])
```
