## Supplementary figures and images for "Pooling size sorted malaise trap fractions to maximise taxon recovery with metabarcoding"

### B_Sequences_merged.pdf

B_merge_PE: Proportion of reads merged using vsearch

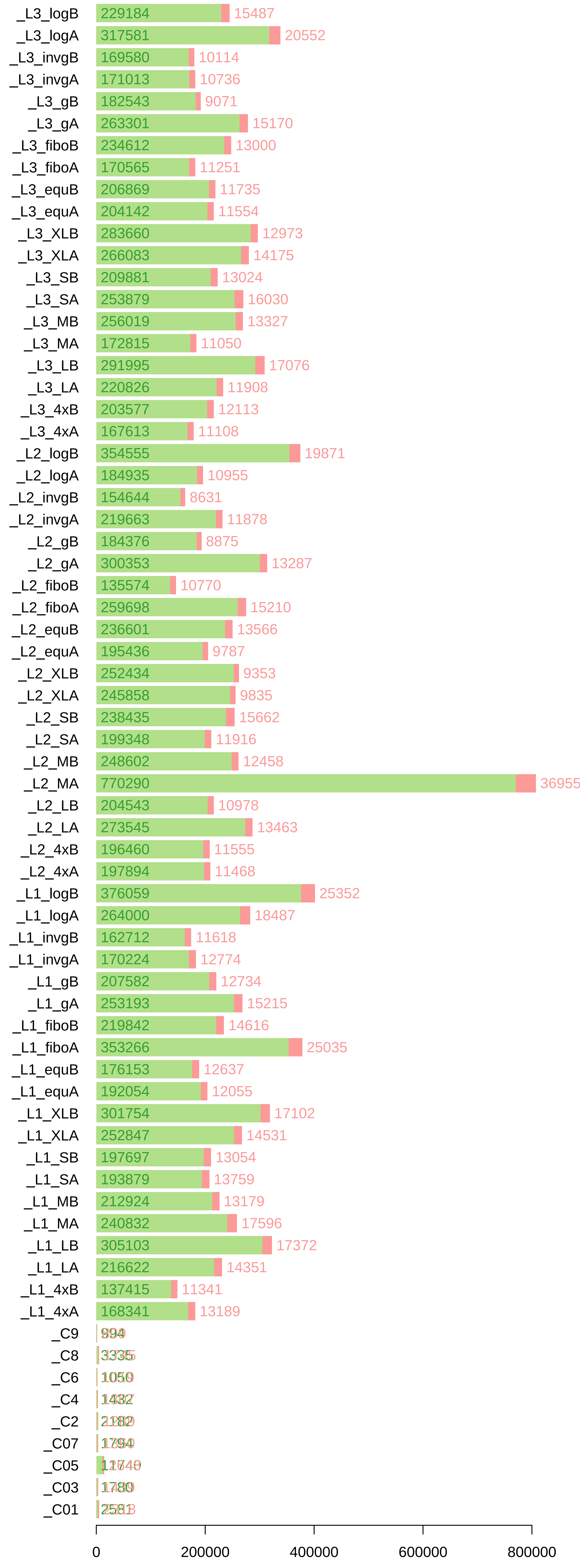

### B_Sequences_merged_rel.pdf

B_merge_PE: Proportion of reads merged using vsearch

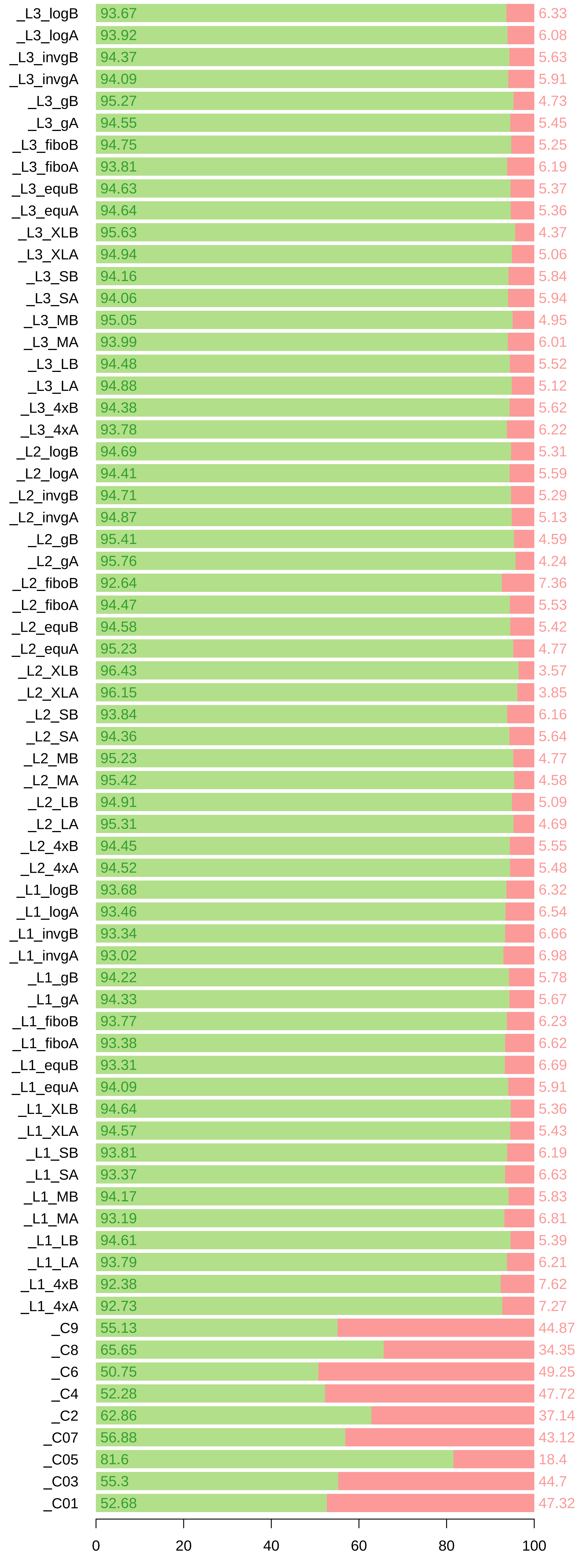

### C_Primers_trimmed.pdf

#### C_Cutadapt: Proportion of reads with no primer detected

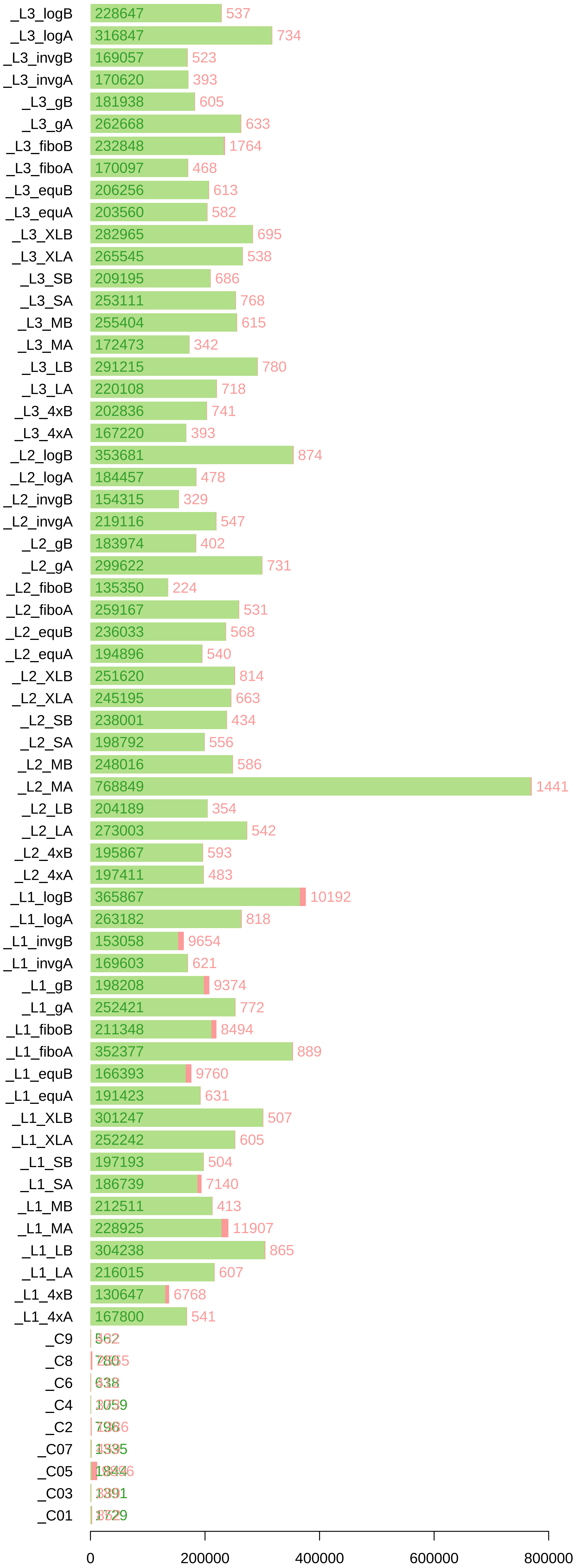

### C_Primers_trimmed_rel.pdf

C_Cutadapt: Proportion of reads with no primer detected

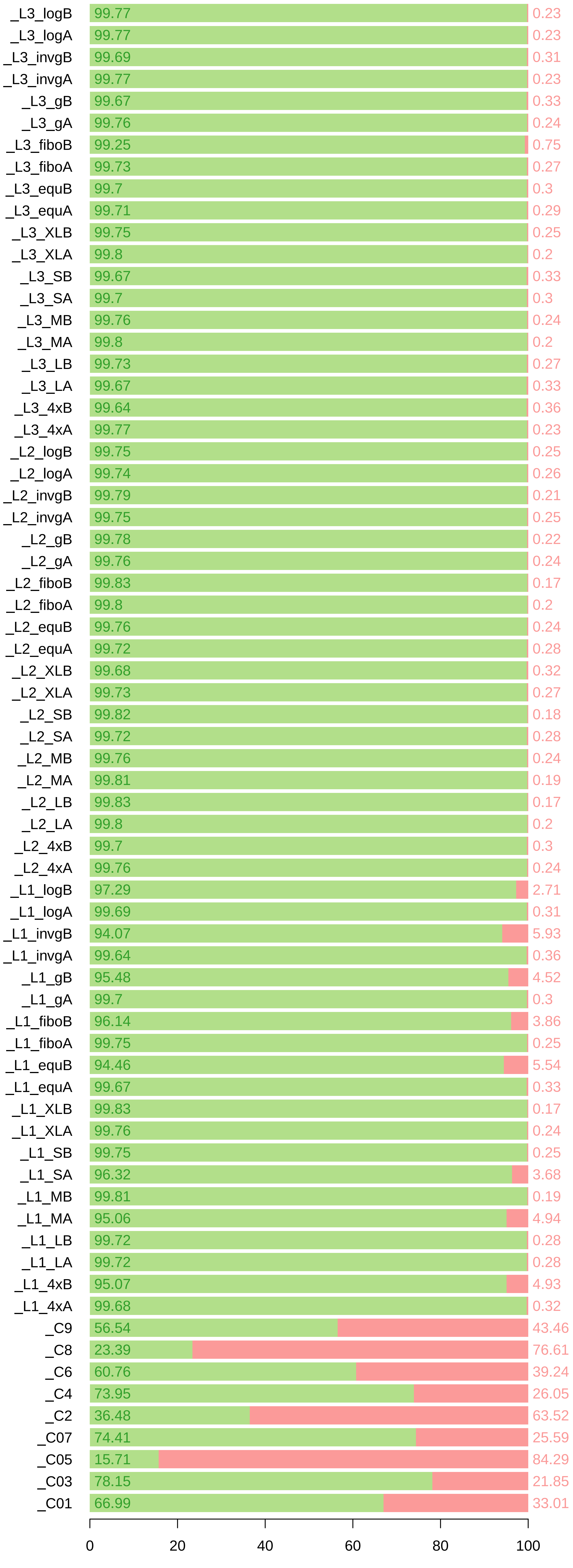

### combined_relSMLXL.pdf

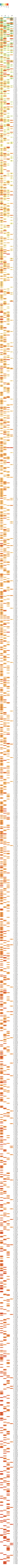

### D_minmax_discarded.pdf

#### D_Minmax: Proportion of reads with 303−323bp length

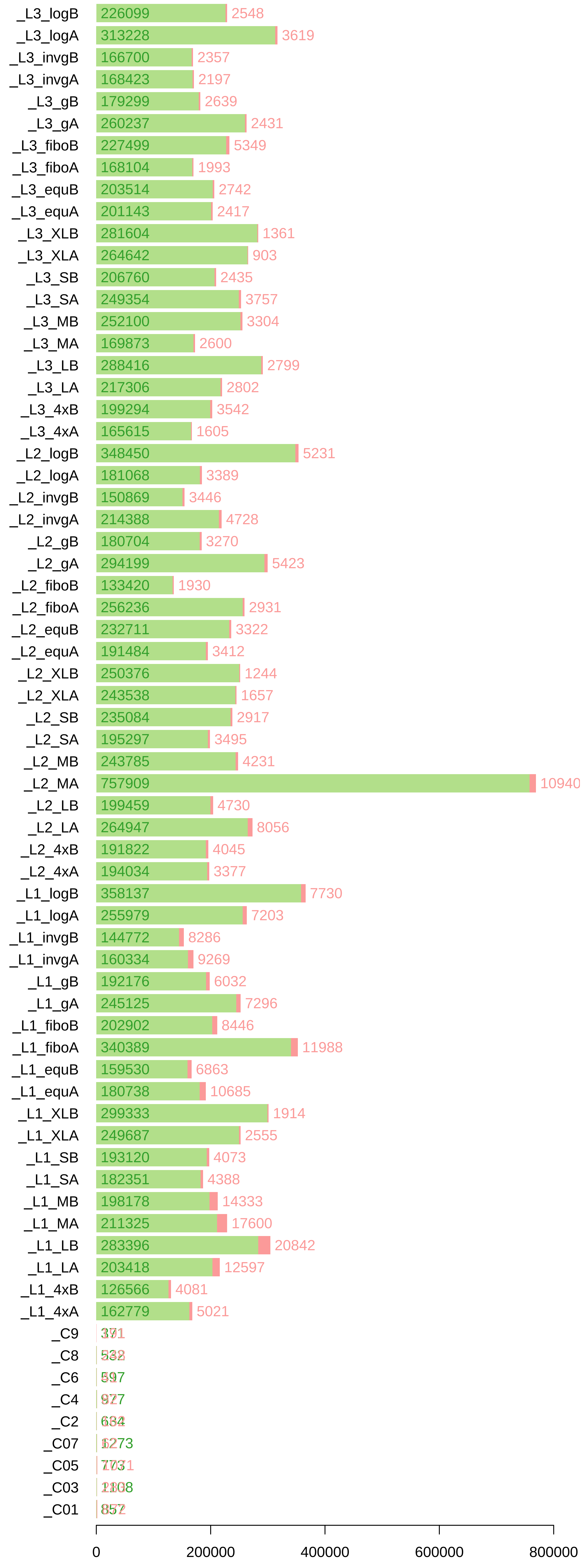

### D_minmax_discarded_rel.pdf

#### D_Minmax: Proportion of reads with 303−323bp length

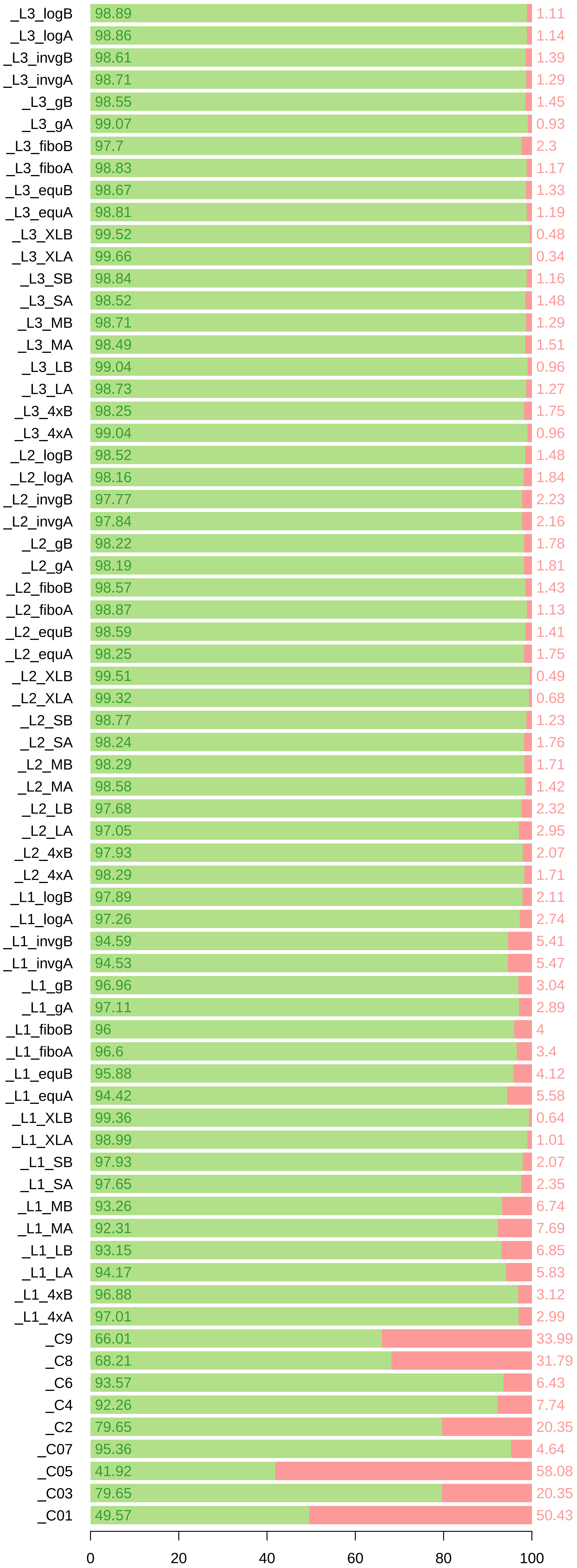

### E_LowQuality_sequences.pdf

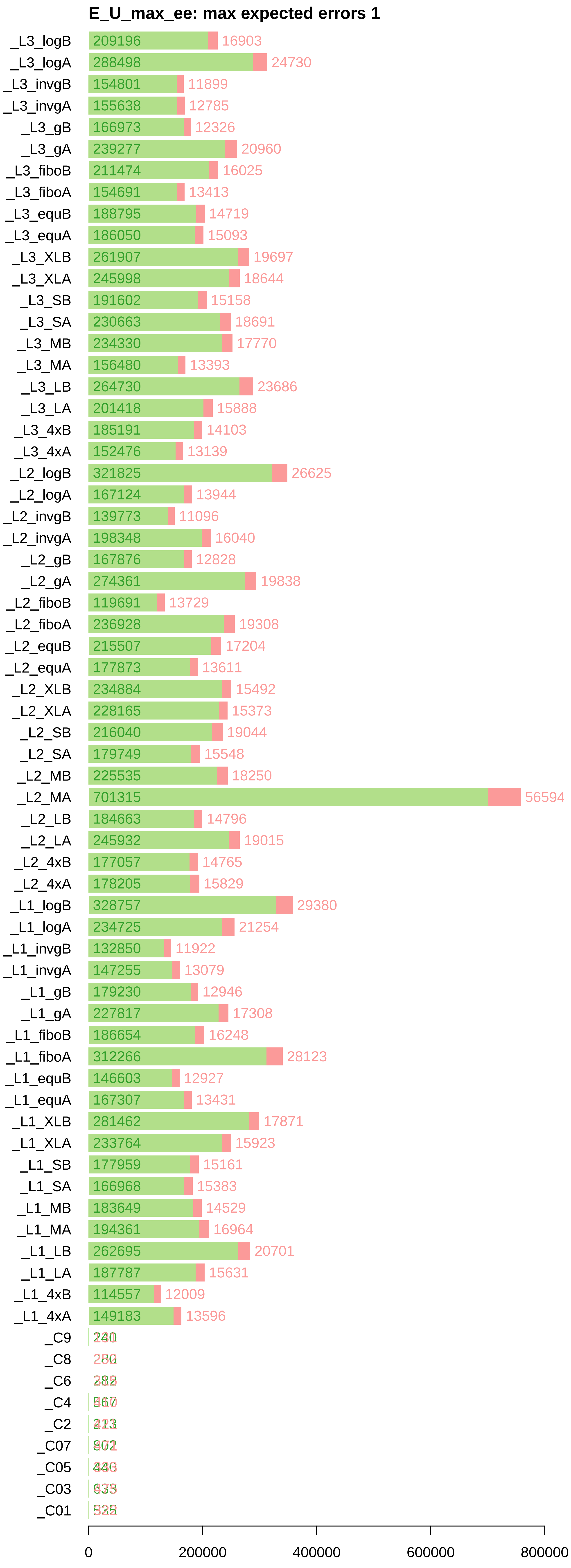

### E_LowQuality_sequences_rel.pdf

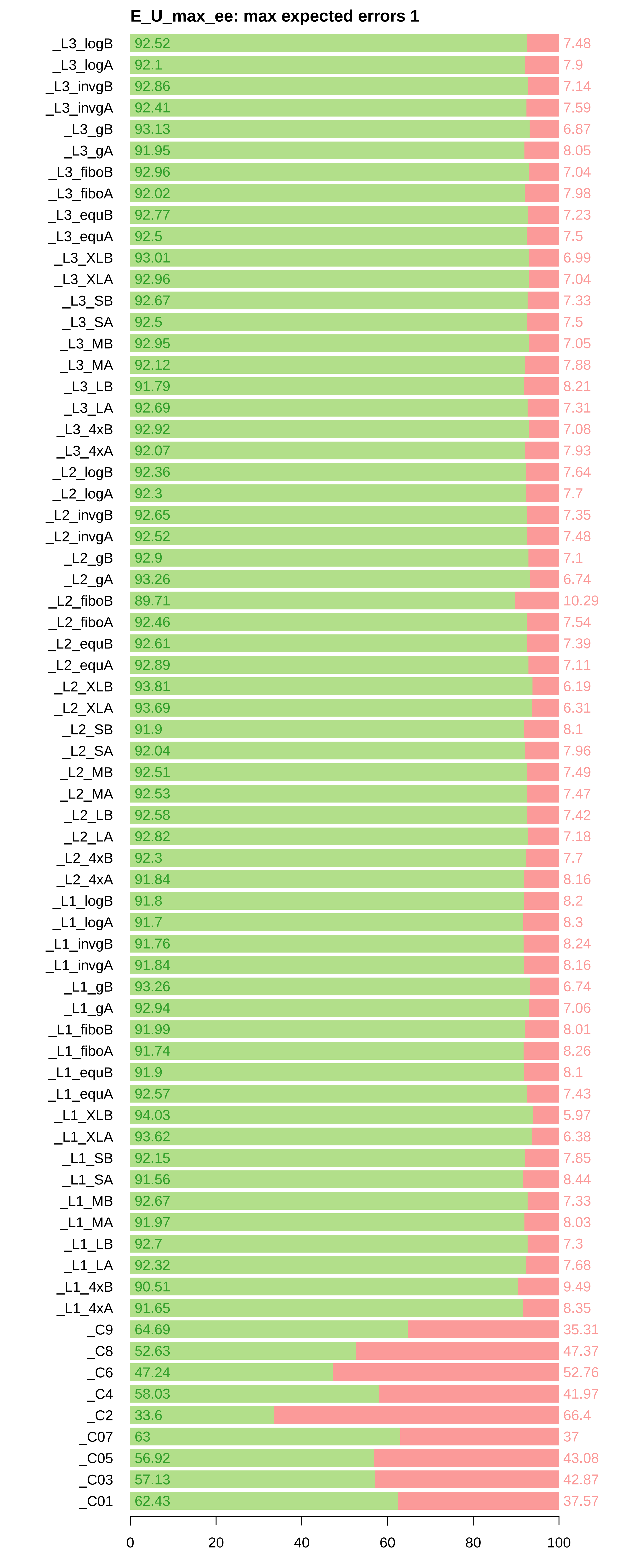
